# Wingless signaling promotes lipid mobilization through signal-induced transcriptional repression

**DOI:** 10.1101/2023.01.25.525602

**Authors:** Mengmeng Liu, Rajitha-Udakara-Sampath Hemba-Waduge, Xiao Li, Xiahe Huang, Tzu-Hao Liu, Xianlin Han, Yingchun Wang, Jun-Yuan Ji

## Abstract

Conserved Wnt/Wingless signaling plays pivotal roles in regulating normal development and energy metabolism in metazoans, and aberrant activation of Wnt signaling drives the pathogenesis of many diseases including cancer. However, the role of Wnt signaling in regulating cellular lipid homeostasis, particularly lipid mobilization, remains poorly understood. Here we show that canonical Wg signaling inhibits lipid accumulation in *Drosophila* larval adipocytes by stimulating lipid catabolism while simultaneously inhibiting lipogenesis. Using a combination of RNA-sequencing and CUT&RUN assays, we identified a battery of Wg target genes encoding key factors required for lipogenesis (such as *FASN1* and *AcCoS*), lipolysis (such as lipid droplet-associated proteins *Lsd-1* and *Lsd-2*), and fatty acid β-oxidation in the mitochondria and peroxisome (e.g., *CPT1* and *CRAT*), most of which are directly repressed by active Wg signaling. Furthermore, lipid accumulation defects caused by active Wg signaling are rescued by either ectopically expressing Lsd-1 and Lsd-2 or depleting the transcriptional repressor Aef1, whose binding motif was identified in 52% of Wg signaling-repressed genes. These findings suggest that active Wg signaling reduces intracellular lipid accumulation by inhibiting lipogenesis and fatty acid β-oxidation and by promoting lipolysis and lipid mobilization, and Wg signaling-induced transcriptional repression play a prominent role in these converging mechanisms.

---

The Wnt/Wingless (Wg) signaling pathway has long been known to play critical roles in regulating development of metazoans, and deregulated Wnt activities are linked to a variety of human cancers, particularly colorectal cancer (Bienz and Clevers, 2000; Cadigan, 2012; Clevers and Nusse, 2012; Fearon, 2011; Nusse and Clevers, 2017; Welters and Kulkarni, 2008). Core components of the canonical Wnt/Wg signaling pathway are evolutionarily conserved in *Drosophila*. Without Wnt/Wg ligand binding, the β-catenin (Armadillo or Arm in *Drosophila*) “destruction complex”, comprised of two scaffold proteins Adenomatosis Polyposis Coli (APC) and Axin (Axn) and two protein kinases Casein-kinase 1α (CK1-α) and Glycogen Synthase Kinase 3 (GSK-3, or Shaggy in *Drosophila*), actively phosphorylates the transcriptional cofactor Arm/β-cat, leading to its degradation in cytoplasm. Consequently, Pangolin (Pan, the TCF/LEF ortholog in flies), the transcription factor downstream of the Wg signaling pathway, is repressed by transcription corepressors such as Groucho (Gro, a *Drosophila* TLE homolog), causing Wnt signaling to be inactive (Bejsovec, 2018; Franz et al., 2017; Nusse and Clevers, 2017). In the presence of Wg ligand binding, the destruction complex can no longer phosphorylate Arm, resulting in Arm accumulation which binds to Pan, thereby activating the expression of Wg target genes (Bejsovec, 2018; Franz et al., 2017; Ramakrishnan et al., 2018).

Compared to its roles in regulating normal development and its dysregulation in human cancers, much less is known about the role of Wnt/Wg signaling in regulating metabolism, particularly lipid metabolism (Abou Azar and Lim, 2021). In mammals, Wnt signaling is known to inhibit adipocyte differentiation, or adipogenesis (Ross et al., 2000; Waki et al., 2007). However, the role of Wnt signaling in regulating lipolysis, β-oxidation of fatty acids (FAs), and lipogenesis is poorly understood. This can be attributed to the complex relationships among adipogenesis, lipogenesis, and lipolysis pathways in mammals which are inextricably intertwined and further complicated by the presence of multiple paralogs that contain partial redundant functions. The fundamental metabolic processes, such as glycolysis, the tricarboxylic acid (TCA) cycle, lipogenesis, lipolysis, and fatty acid β-oxidation (FAO), are virtually identical in vertebrates but simpler in invertebrates such as *Drosophila* (Arrese and Soulages, 2010; Heier and Kuhnlein, 2018). Interestingly, adipogenesis in *Drosophila* occurs only during late embryonic stage, temporally separated from lipogenesis, lipolysis, and FAO that occur during postembryonic stages. Thus, *Drosophila* provides an ideal system to elucidate the role of Wnt signaling in regulating these different aspects of lipid homeostasis.

## Wg signaling promotes lipid mobilization and reduces fat accumulation

We have previously reported that Wg signaling is hyperactive in *Axn^127^* homozygous mutant larvae, resulting in defective fat accumulation (Zhang et al., 2017). To study how active Wg signaling reduces fat accumulation, we used a transgenic RNA interference (RNAi) approach to deplete either Axn or Slmb (Supernumerary limbs), a conserved E3 ubiquitin ligase required for the degradation of Arm (Jiang and Struhl, 1998). As expected, silencing *Axn* or *slmb* in larval adipocytes significantly increased the levels of Arm (Suppl Fig. 1a-f), which correlated to elevated expression of the *fz3-RFP* (Fig. 1a-c) and *nkd^EGFP^* reporters (Suppl Fig. 1g,h), which are both expressed under the direct control of Wg signaling in the *Drosophila* gut (Tian et al., 2017). Interestingly, the size of adipocytes appeared heterogeneous, with most small adipocytes mixed among occasional large adipocytes (Fig. 1b-c, 1e-f). Smaller adipocytes had high levels of nuclear Arm and elevated expression of *fz3-RFP* and *nkd^EGFP^* (Fig. 1b-c; Suppl Fig. 1a-h), suggesting that Wg signaling was more active in small adipocytes than in the adjacent large adipocytes. The underlying mechanism of this larval adipocyte heterogeneity is currently unknown. Importantly, depleting either *Axn* or *slmb* using multiple adipocyte-specific Gal4 lines reduced the size of adipocytes, the number of lipid droplets, and triglycerides (TGs) (Fig. 1b,c, 1e-f; Suppl Fig. 1i-m), which could be rescued by depletion of Pan (Suppl Fig. 1n-o). Thus, Wg signaling is active in small adipocytes with reduced fat accumulation, in agreement with the effects of canonical Wg signaling in disrupting fat accumulation in *Drosophila* larvae (Zhang et al., 2017).

**Fig. 1.**
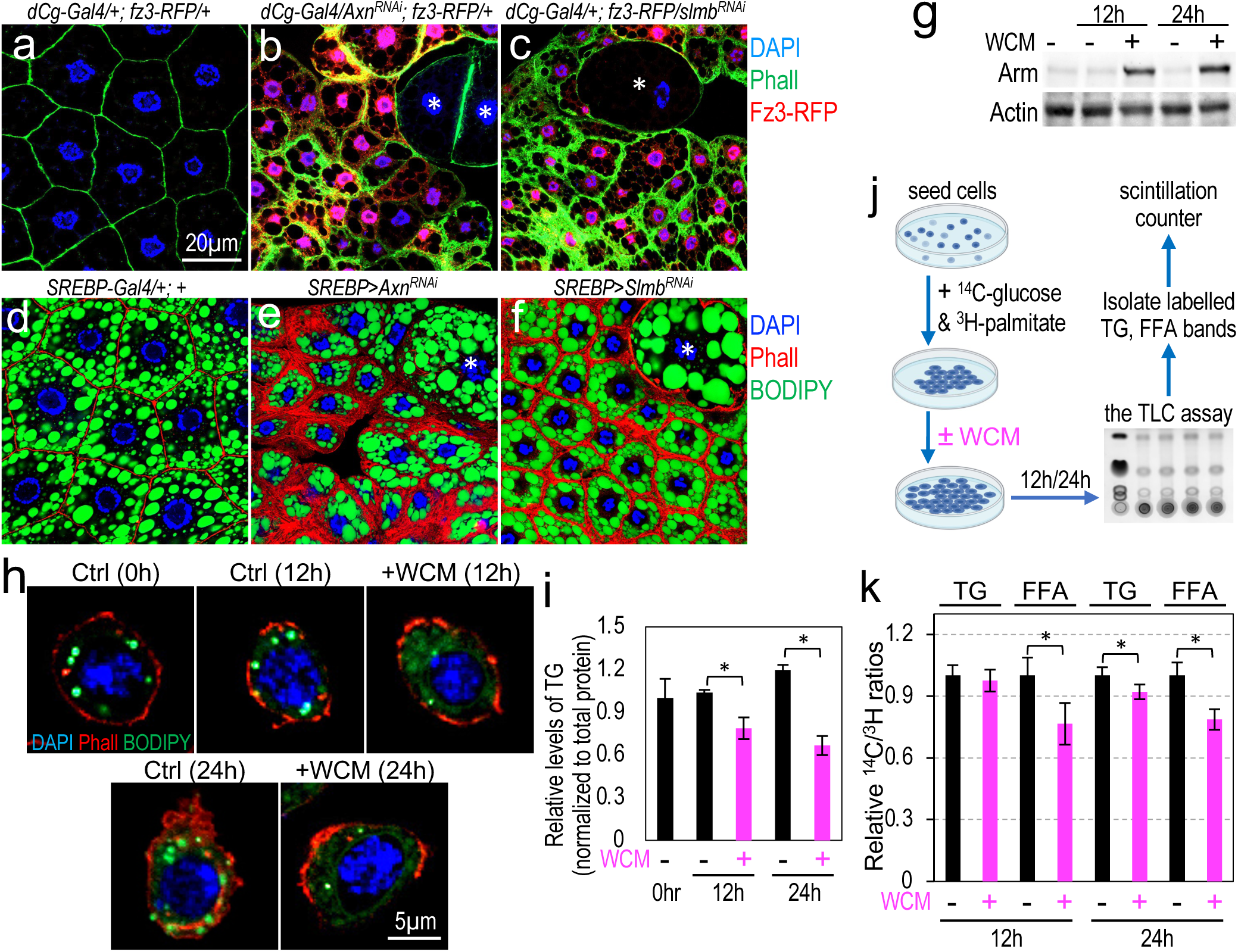
Active Wnt signaling disrupts lipid homeostasis in larval adipocytes. (a-c) Representative confocal images of larval adipocytes from larvae of indicated genotypes stained with DAPI (blue) and BODIPY (green). Red: *fz3*-driven expression of RFP. Note the elevated expression of *fz3-RFP* in small adipocytes in b/c, in contrast to low or no expression of *Fz3-RFP* in control (a) and the large adipocytes (* in b/c). Genotypes: (a) *dCg-Gal4/+; fz3-RFP/+*; (b) *dCg-Gal4/UAS-Axn^RNAi^; fz3-RFP*/+; and (c) *dCg-Gal4/+; fz3-RFP*/*UAS-slmb^RNAi^.* (d-f) Confocal images of larval adipocytes of the indicated genotypes stained with DAPI (blue; nuclei), Phall (Phalloidin, red; microfilament bundles under cell membrane), and BODIPY (green; lipid droplets). Large adipocytes are marked with asterisks (* in e/f). Genotypes: (d) *SREBP-Gal4/+; +*; (e) *SREBP-Gal4/UAS-Axn^RNAi^; +*; (f) *SREBP-Gal4/+; UAS-slmb^RNAi^/+; +.* (g) Western blot analysis of Arm using S2R+ cells treated with WCM (Wg-conditioned medium). Actin was used as a loading control. (h) Representative confocal images of cultured S2R+ cells stained with DAPI (blue), Phall (red), and BODIPY (green). Scale bar: 5μm. (i) Quantification of TG levels, and the results are normalized to the control (n = 3, independent biological repeats). (j) The experimental design of the dual isotope radiolabeling experiments using cultured S2R+ cells. TLC: thin layer chromatography. (k) Relative ^14^C/^3^H ratios in TG and FFAs of S2R+ cells treated with WCM for 12h or 24h.

To test whether reduced lipid accumulation in larval adipocytes was due to increased lipolysis or reduced lipogenesis, we used dual radioisotope labeling using ^14^C-labeled glucose and ^3^H-palmitic acid to directly assess the effects of Wg signaling on the breakdown and synthesis of free fatty acids (FFAs) and TGs. Through glycolysis, ^14^C-glucose is converted into acetyl-CoA, which can be oxidized to CO_2_ through the tricarboxylic acid cycle or converted into FFAs and TGs via lipogenesis; ^3^H-palmitic acid can be either directly esterified with glycerol (may be ^14^C-labeled) to produce TGs or oxidized via β-oxidation to generate acetyl-CoA, which can be further oxidized into CO_2_ (Suppl Fig. 2a). Thus, the rate of lipolysis or lipogenesis can be measured by the relative rates of ^14^C and ^3^H accumulation (the ^14^C/^3^H ratio) in TGs and FFAs, where a reduced ^14^C/^3^H ratio indicates predominantly increased lipolysis and vice versa (Suppl Fig. 2b) (Gibbons et al., 1994; Huynh et al., 2014; Leibel, 1985; Leibel et al., 1984). To avoid compounding factors such as the presence of heterogeneous cell populations in larvae fed with fly food containing carbohydrate-rich and imprecisely defined ingredients, cultured *Drosophila* S2 cells, known to accumulate oil droplets (Beller et al., 2006; Guo et al., 2008; Walther and Farese, 2012), were used. For these experiments, we used the fact that depletion of Axn in S2 cells activates Wg signaling (Gonsalves et al., 2011), as does treating cultured *Drosophila* cell lines such as Kc167 cells, S2R+ cells (Fig. 1g), or cl-8 cells with Wg-conditioned medium (WCM) (Franz et al., 2017; van Leeuwen et al., 1994; Yanagawa et al., 1998); WCM treatment also reduced the number of lipid droplets and the level of TGs (Fig. 1h, i). These results indicate that the effect of active Wg signaling in reducing TG accumulation is not limited to larval adipocytes. ^14^C-labeled glucose and ^3^H-palmitic acid were added to the medium and then treated the S2R+ cells with WCM to activate Wg signaling. TGs and FFAs were separated with thin-layer chromatography (TLC) and ^14^C/^3^H ratios were quantified using liquid scintillation (Fig. 1j). This analysis revealed that the ^14^C/^3^H ratios of both FFA and TGs were significantly reduced in S2R+ cells after WCM treatment (Fig. 1k), which constitutes direct evidence for the role of Wg signaling in stimulating lipolysis and inhibiting *de novo* lipogenesis.

## Wnt signaling represses the transcription of genes encoding factors that control lipid catabolism and *de novo* lipogenesis

Our previous analyses of the *Axn^127^* homozygous mutant larvae revealed that hyperactive Wg signaling caused a general reduction of most types of TGs, but increases the levels of various FFAs (Zhang et al., 2017). Those effects were validated in larvae with adipose tissue-specific activation of Wg signaling (Fig. 2a). To gain insights into the molecular mechanism behind the Wnt signaling-stimulated lipid catabolism and reduction of lipogenesis, we carried out bulk RNA sequencing (RNA-seq) analyses of dissected larval fat body with specific depletions of *Axn* or *slmb*. A total of 1622 upregulated genes and 1780 downregulated genes were identified in both *Axn^RNAi^* and *slmb^RNAi^* adipocytes, compared to the controls (Fig. 2b; both adjusted p- and q-values were less than 0.05). As expected, “Pathway” and “Gene Ontology” cluster analyses (Boyle et al., 2004) revealed that the ‘Wnt signaling pathway’ was significantly activated in both *Axn^RNAi^* and *slmb^RNAi^* fat bodies (Fig. 2c), and upregulated genes in this pathway included direct Wg target genes such as *fz3* (*frizzled 3*), *nkd* (*naked cuticle*), and *Notum* (Suppl Fig. S3a). Multiple biological pathways were also significantly upregulated and downregulated by active Wg signaling (Fig. 2c, Suppl Fig. S3b,c). Notably, several lipid metabolism-related pathways were significantly downregulated in adipocytes with active Wg signaling, including ‘FA degradation,’ ‘FA biosynthesis,’ ‘FA metabolism,’ ‘oxidative phosphorylation,’ ‘TCA cycle,’, and ‘peroxisome’ pathways (Fig. 2c).

**Fig. 2.**
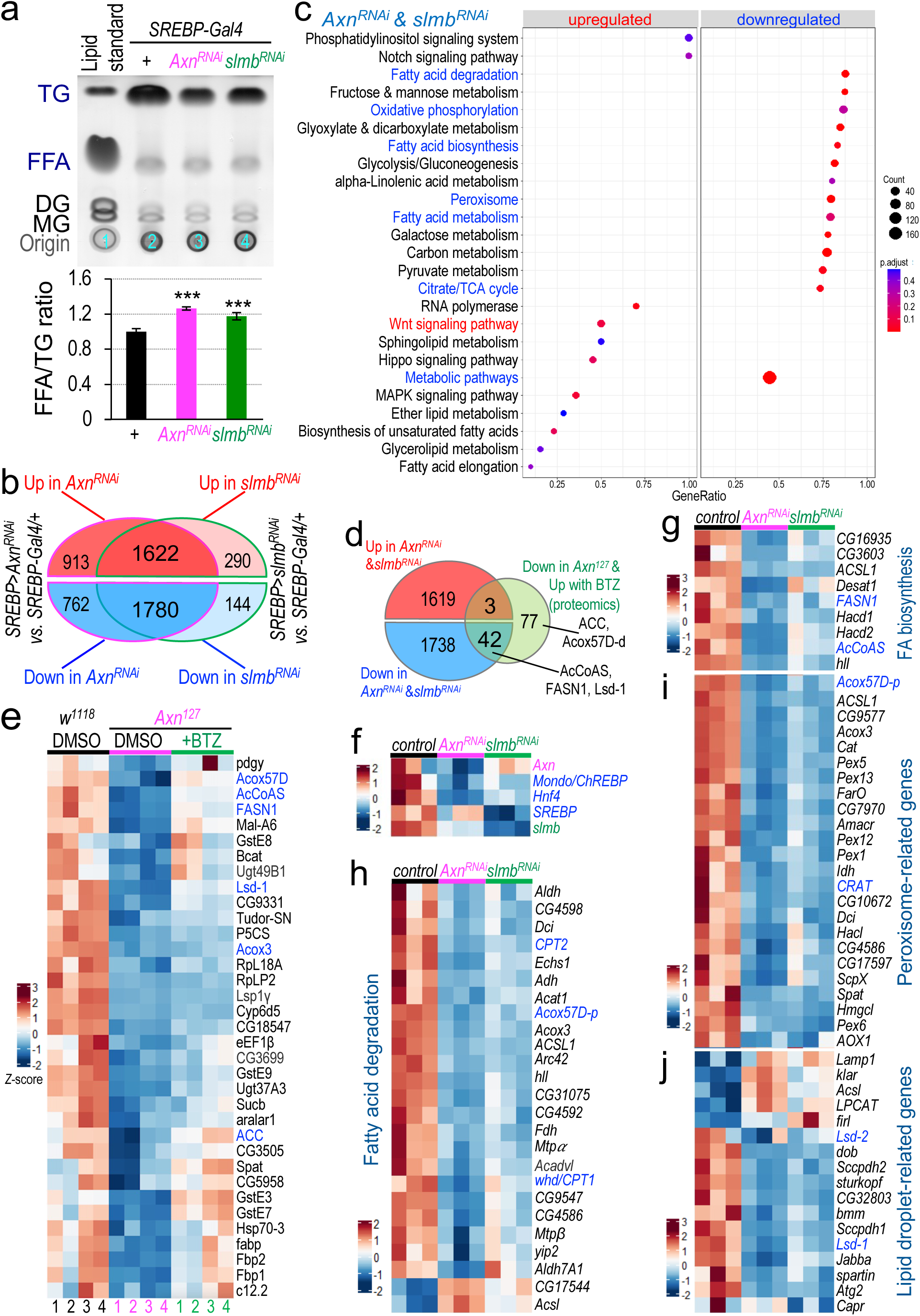
Wnt signaling represses the transcription of genes encoding factors that control *de novo* lipogenesis and lipid catabolism. (a) Analyses of the relative levels of FFAs and TGs using the TLC assay in larvae with fat body-specific depletion of either *Axn* or *slmb*. Genotypes: *SREBP-Gal4/+; +* (lane #2); *SREBP-Gal4/UAS-Axn^RNAi^; +* (lane #3); and *SREBP-Gal4/+; UAS-slmb^RNAi^/+* (lane #4). Lower panel: quantification of the FFA/TG ratios. (b) Venn diagram showing the number of genes altered in fat body with the depletion of either *Axn* or *slmb*. (c) Common pathways related to metabolism that are significantly altered in larval fat body with the depletion of either *Axn* or *slmb*. Downregulated pathways related to carbohydrate and lipid metabolism are highlighted in blue; as expected, Wnt signaling pathway (highlighted in red) is significantly upregulated. (d) Venn diagram showing the number of genes altered in *Axn^RNAi^* or *slmb^RNAi^* fat body, as compared to the number of proteins that are downregulated in *Axin^127^* mutant larvae but reversed by feeding the larvae with 2μM of BTZ in fly food. (e) Heatmap showing the levels proteins that are reduced in *Axin^127^* mutant larvae but reversed by BTZ as quantified by proteomics. Key proteins such as ACC, AcCoAS, Acox-57D, Acox3, FASN1, and Lsd-1 are highlighted in blue. (f) Heatmap of the mRNA levels of *Mondo/ChREBP*, *Hnf4* and *SREBP* in *Axn^RNAi^* or *slmb^RNAi^* fat body. The mRNA levels of *Axn* and *slmb* serve as positive controls. (g) Heatmap of gene expression for genes of the ‘fatty acid biosynthesis’ pathway. (h) Heatmap of the genes in the ‘fatty acid degradation’ pathway. (i) Heatmap of the genes in the ‘peroxisome’ pathway. (j) Heatmap of the genes in the lipid droplet-related genes. Data shown in these heatmaps represent z-score normalized average normalized count values from triplicates per genotype. Key genes that were further analyzed are highlighted in blue.

Given that the absolute steady-state levels of mRNAs do not always strictly correlate with the cellular abundance of their corresponding proteins (Schwanhausser et al., 2011), we performed quantitative proteomic analyses with TMT (tandem mass tag) labeling (Bi et al., 2014; Unwin, 2010; Wiese et al., 2007). Feeding the *Axn^127^* mutant larvae with peptide boronic acids such as Bortezomib (BTZ) rescued the defects in fat accumulation (Zhang et al., 2017), thus we analyzed *Axn^127^* mutant larvae and the effects of BTZ treatment on protein levels. Next, we compared the list of proteins to the list of genes whose mRNA levels were significantly altered in *Axn^RNAi^* and *slmb^RNAi^* fat bodies. This led to the identification of 45 proteins whose levels were changed in the same manner as their mRNA levels (Fig. 2d). Of these 45 proteins, 42 proteins were significantly lowered in *Axn^127^* mutants and upregulated by BTZ (Fig. 2d). Five of these 42 proteins were particularly intriguing including FASN1 (FA synthase 1) and ACC (acetyl-CoA carboxylase), the key enzymes required for triglyceride biosynthesis; AcCoAS (acetyl Coenzyme A synthase, which converts acetate into acetyl-CoA, a central metabolic intermediate linking FA synthesis and the TCA cycle); acyl-CoA oxidase Acox57D that functions in peroxisomal FAO; and Lsd-1 (Lipid storage droplet-1, the Perilipin-1 homolog in *Drosophila*) (Fig. 2e). These results indicate that hyperactive Wg signaling regulates intracellular lipid homeostasis through certain key factors involved in FA synthesis, FAO, and lipid droplet-associated proteins (LDAPs) on lipid droplets (LDs).

As quantitative proteomic analyses may miss the detection of proteins with low abundance in larvae, we analyzed the RNA-seq data by focusing on genes involved in FA biosynthesis, FAO, LDAPs, and peroxisome-related genes. This analysis revealed a general reduction of gene encoding enzymes and factors involved in FA biosynthesis and degradation by active Wg signaling (Fig. 2f-h, Suppl Fig. S3d). Given the critical roles of SREBP and Mondo/ChREBP in regulating the transcription of lipogenic enzymes, and the role of dHnf4 in regulating the transcription of factors required for FAO, it is interesting that the mRNA levels of these key transcription factors were also significantly reduced in *Axn^RNAi^* and *slmb^RNAi^* fat bodies (Fig. 2f). Reduction of SREBP and Mondo/ChREBP is consistent to the reduced expression of their target genes such as *FASN1*, *AcCoAS*, and *ACSL1* (long-chain fatty acid-CoA ligase) (Fig. 2g). Similarly, reduced *dHnf4* mRNA level also correlated to a significant reduction of *whd* (*withered*, encoding carnitine O-palmitoyltransferase CPT1), *CPT2*, *CG4598* and *yip2* (Fig. 2h); it has been reported that the transcription of these genes is regulated by Hnf4 in *Drosophila* (Palanker et al., 2009). CPT1 and CPT2 are required for transporting fatty acids into the mitochondria for FAO, while *CG4598* and *yip2* encode enzymes involved in FAO in the mitochondria. In addition to the mitochondria, peroxisomes also play critical roles in β-oxidatin of FAs (Reddy and Hashimoto, 2001; Schrader et al., 2015). Active Wg signaling also reduced the expression of *CRAT*, encoding carnitine O-acetyl-transferases that transport very long chain FAs into peroxisomes, as well as multiple genes encoding Peroxins and other peroxisomal proteins (Fig. 2i, Suppl Fig. S3e). Downregulation of these key factors involved in FAO in *Axn^RNAi^* and *slmb^RNAi^* fat bodies suggests that Wg signaling negatively regulates FAO in peroxisomes and mitochondria. Consistent to the results of the dual radioisotope labeling assay in S2R+ cells (Fig. 1k), these observations in larval adipocytes suggest that active Wg signaling inhibits both *de novo* lipogenesis and FAO.

Identification of Lsd-1 in proteomic analyses prompted us to extend this analysis to other LDAPs. Besides *Lsd-1*, a few genes encoding LDAPs such as *Lsd-2*, *Jabba*, *spartin*, and *sturkopf* were also significantly reduced in *Axn^RNAi^* and *slmb^RNAi^* fat bodies (Fig. 2j). *Drosophila* has two perilipin orthologs, Lsd-1 (Perilipin-1) and Lsd-2 (Perilipin-2), which impede lipolysis and lipid mobilization by restraining lipases from accessing TGs within lipid droplets (Tirinato *et al*., 2017). Thus, downregulation of LDAP expression in adipocytes by active Wg signaling may explain how Wg signaling promotes lipolysis.

To validate the repressive effects of Wg signaling on the transcription of *SREBP*, *Hnf4*, *Lsd1*, *Lsd2*, *Acox57D-d*, *FASN1*, *AcCoAS*, and *ACC* at the cellular level, their mRNA transcripts were analyzed using the multiplexed *in situ* hybridization chain reaction (HCR) RNA fluorescence *in situ* hybridization (RNA-FISH) imaging technology (Choi et al., 2018). Results from this these analyses confirmed that mRNA levels of *Hnf4* (Fig. 3a-c), *Lsd1*, *Lsd2*, and *Acox57D-d* (Fig. 3d-f), as well as *FASN1*, *AcCoAS*, and *ACC* (Fig. 3g-j) were significantly reduced in small adipocytes when compared to the controls or the adjacent large adipocytes in *Axn^RNAi^* and *slmb^RNAi^* fat bodies (Fig. 3a, 3d, 3g). The effect of Wg signaling on mRNA levels of *SREBP* was less conclusive (Fig. 3a-c). Elevated expression of *arm* was evident in small adipocytes (Fig. 3a-c), consistent with more active Wg signaling in small adipocytes (Fig. 1a-f). Together with the RNA-seq and quantitative proteomic analyses (Fig. 2e-i), these results show that the transcription of *Hnf4*, *Lsd1*, *Lsd2*, *Acox57D-d*, *FASN1*, *AcCoAS*, and *ACC* is repressed by active Wg signaling in adipocytes.

**Fig. 3.**
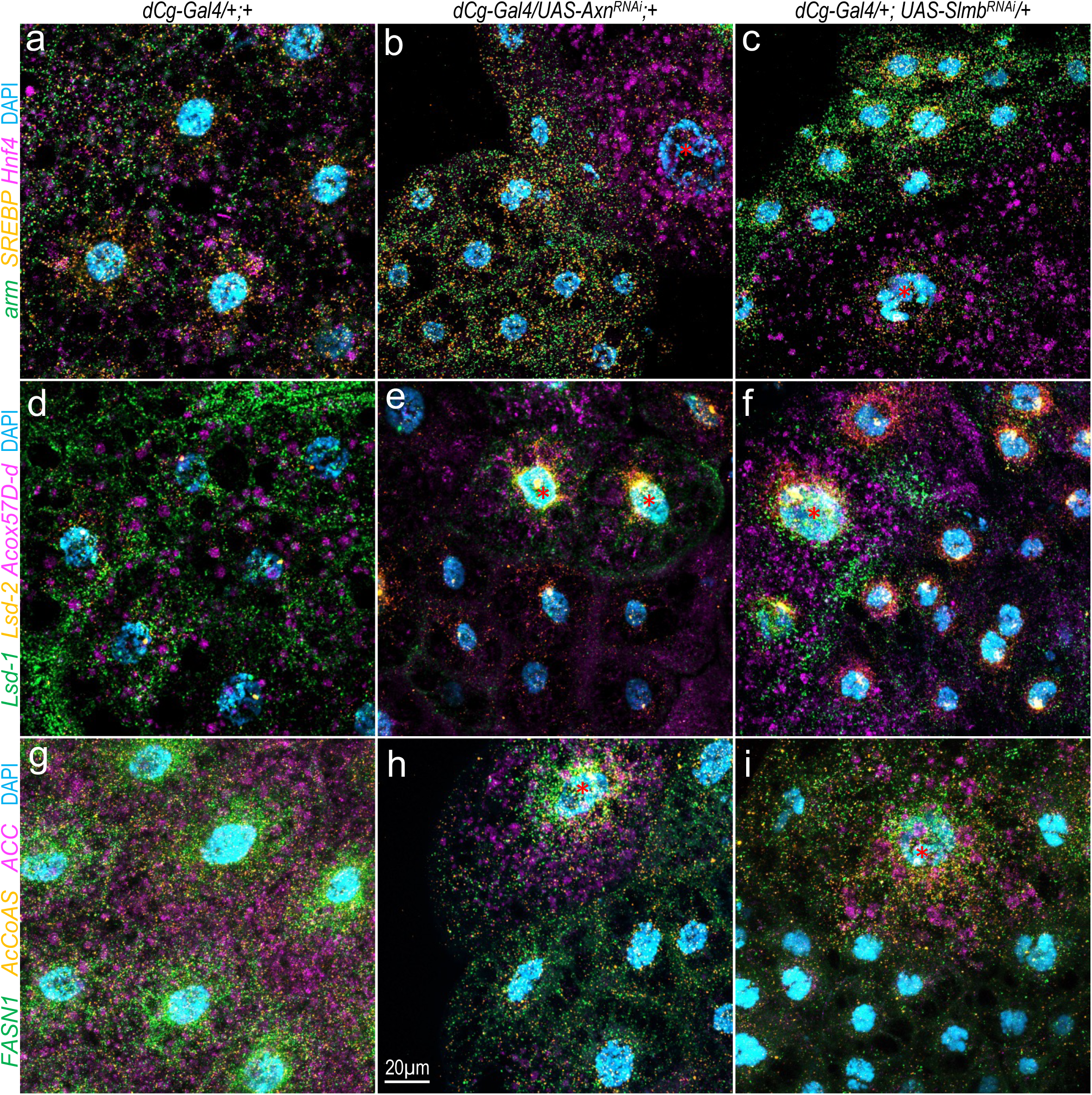
Wg signaling regulates the transcription of genes encoding factors that control intracellular lipid homeostasis. (a-c) The mRNA transcripts of *arm* (green), *Hnf4* (magenta) and *SREBP* (orange) in larval adipocytes detected by the HCR assay. Note the higher levels of *Hnf4* and *SREBP* in big adipocytes than the neighboring small adipocytes, which have higher levels of *arm* as expected. (d-f) The mRNA transcripts of *Lsd-1* (green), *Acox57D-d* (magenta) and *Lsd-2* (orange) in larval adipocytes, note the lower levels of the transcripts of these genes in small adipocytes than the bigger adipocytes. (g-i) Lower level of FASN1 (green), AcCoAS (orange) and ACC (magenta) in small adipocytes. The big adipocytes with low Wg activity are indicated with asterisks (red). The mRNA transcripts of these genes were detected by *in situ* hybridization chain reaction (HCR) assay. The probe sets that these genes were used with are B1-Alexa Fluor 488 (green), B2-Alexa Fluor 594 (orange) and B3-Alexa Fluor 647 (magenta) amplifiers, respectively, in multiplexed in situ HCR. These images are projections of 12-15 successive optical sections (1 μm apart).Genotypes: (a) *dCg-Gal4/+; +*; (b) *dCg-Gal4/UASAxn^RNAi^; +*; and (c) *dCg-Gal4/+; UAS-slmb^RNAi^/+*. Scale bar: 20 μm.

Although it is unknown whether these inhibitory effects are conserved in other species, the inhibitory effects of Wnt signaling on *FAS* gene expression and lipogenesis have been reported in mouse adipocytes and the liver of juvenile turbot treated with GSK3 inhibitor LiCl (Bagchi et al., 2020; Liu et al., 2016). Much less is known about potential role of Wnt signaling in regulating lipolysis. Hence, we focused our further analyses on the role of Wnt/Wg signaling in lipolysis and lipid mobilization.

## Wg signaling stimulates lipolysis through LDAPs

Identification of Wg signaling-induced repression of *Lsd-1* and *Lsd-2* prompted us to hypothesize that activate Wg signaling reduces LDAPs (e.g., Lsd1 and Lsd2) on the surface of lipid droplets in adipocytes. This could allow for triglycerides within those droplets increased access to lipases, thus favoring the lipid mobilization from tri- to di-to mono-glycerides then to FAs. To test whether reducing *Lsd-1* could enhance the defective fat accumulation driven by Wg signaling, *Lsd-1* was depleted separately or in combination with *Axn* in larval adipocytes. Analyses of lipid droplet morphology and lipid metabolism were then conducted. In agreement with the observation that reducing Lsd-1 enlarges the size of lipid droplets (Beller et al., 2010), depleting *Lsd1* resulted in significantly larger but fewer lipid droplets per adipocyte (Fig. 4c cf. 4a), as well as reduced TG accumulation (Fig. 4g). These observations suggest that lipid droplets coated with less Lsd-1 are prone to fuse with each other and form large lipid droplets and favor lipolysis. Additionally, depleting both *Lsd1* and *Axn* leads to even bigger lipid droplets (Fig. 4d cf. 4c), as well as a further reduction of TG accumulation (Fig. 4g). If Wg signaling represses *Lsd1* transcription, then reducing Wnt signaling is expected to rescue the LD phenotype caused by depleting *Lsd1*. Indeed, simultaneous depletion of *Arm* and *Lsd-1* in the larval adipocytes significantly rescued the large LD phenotype caused by depleting *Lsd-1* (Fig. 4f cf. 4c; 4g), supporting a repressive effect of Wg signaling on *Lsd-1* transcription.

**Fig. 4.**
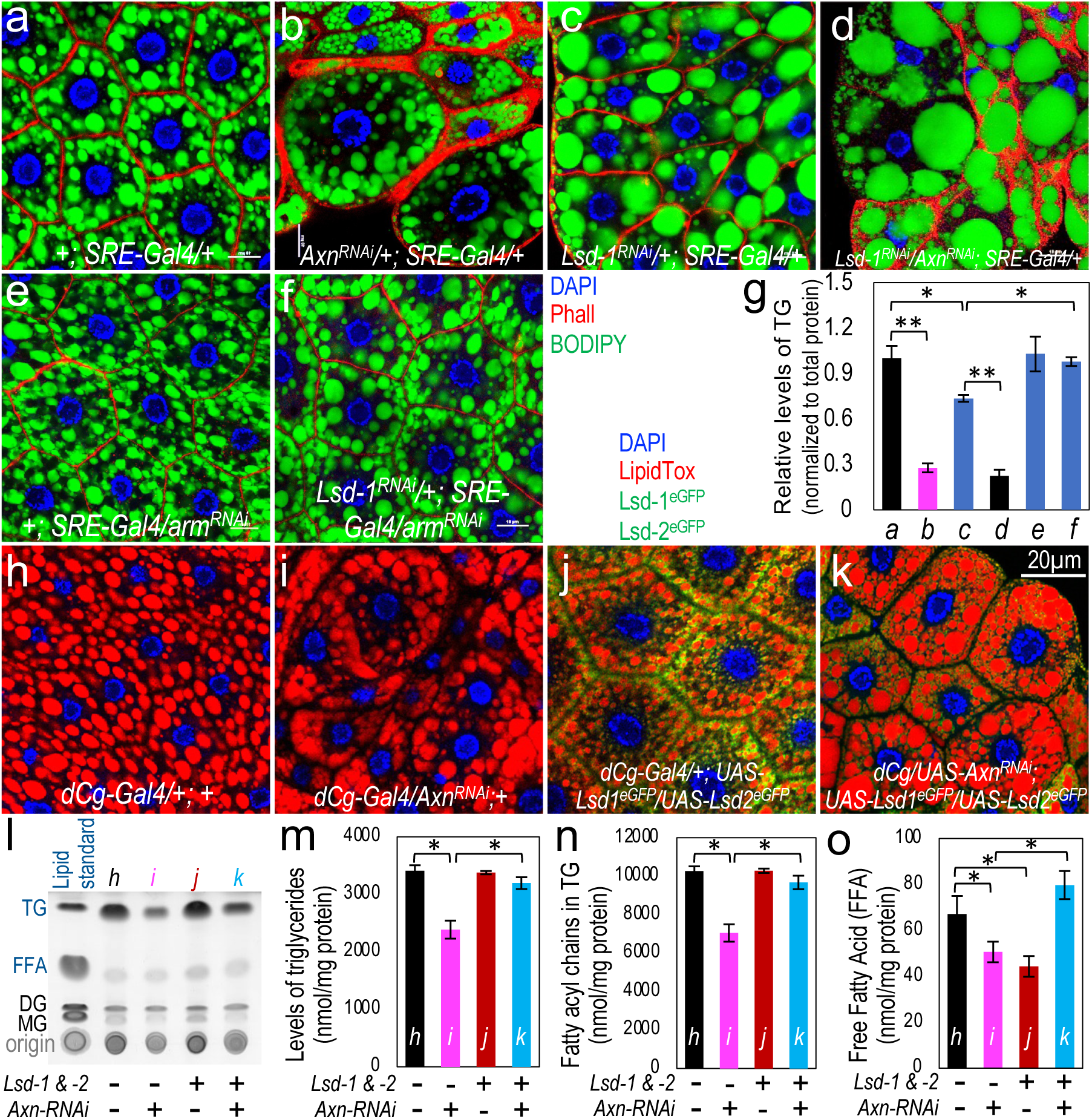
Interplay between lipid droplet-associated proteins and Wnt signaling. (a-f) Representative confocal images of larval adipocytes of the indicated genotypes stained with DAPI (blue), Phall (red), and BODIPY (green). Genotypes: (a) *+; SREBP-Gal4/+*; (b) *UAS-Axn^RNAi^/+; SREBP-Gal4/+*; (c) *UAS-Lsd1^RNAi^/+; SREBP-Gal4/+*; (d) *UAS-Axn^RNAi^/UAS-Lsd1^RNAi^; SREBP-Gal4/+*; (e) *+; SREBP-Gal4/UAS-arm^RNAi^*; and (f) *UAS-Lsd1^RNAi^/+; SREBP-Gal4/UAS-arm^RNAi^*. (g) Quantification of TG levels of the indicated genotypes (n = 3 biological repeats). Results are normalized to the control (*+; SREBP-Gal4/+*). (h-k) Confocal images of larval adipocytes of the indicated genotypes stained with DAPI (blue) and LipidTox (red). Genotypes: (h) *dCg-Gal4/+; +*; (i) *dCg-Gal4/UAS-Axn^RNAi^; +*; (j) *dCg-Gal4/+; UAS-plin1^eGFP^/UAS-Lsd2^eGFP^*; and (k) *dCg-Gal4/UAS-Axn^RNAi^; UAS-Lsd1^eGFP^/UAS-Lsd2^eGFP^*. (l) Chromatographic separation of lipids from larvae of the indicated genotypes using the TLC analysis. Lane 1 shows the lipid standard, a mixture of monoglyceride (MG), diglyceride (DG), TG, and stearic acid (representing FFAs). (m-o) Quantitative lipidomics measurement of TGs (m), Fatty acyl chains in TGs (the amount of each fatty acyl chain in total TG pool; n), and FFAs (o) in fat bodies dissected from the third instar larvae of the indicated genotypes.

Next, we investigated whether ectopic expression of LDAPs could at least partially alleviate the effects of Wg signaling on lipid mobilization. Given that Lsd-1 and Lsd-2 display partially redundant roles in lipid metabolism (Beller et al., 2010; Bi et al., 2012), we tested whether gain of Lsd-1, Lsd-2, or both, could rescue adipocyte defects caused by active Wg signaling. While ectopic expression of either *Lsd-1* or Lsd-2 alone partially rescued the adipocyte defects caused by depletion of *Axn* in the fat bodies (Suppl Fig. S4), ectopic expression of both Lsd-1 and Lsd-2 strongly rescued the fat-body defects caused by Wg signaling (Fig. 4k cf. 4i, controls are shown in Fig. 4h and 4j). Quantification of these effects using TLC showed that ectopic expression of *Lsd-1* and *Lsd-2* strongly rescued the effects of Axn depletion on TG levels (Fig. 4i, comparing the TG levels shown in lane *k* versus lane *i*).

To precisely measure these effects on different types of TGs and FFAs, larval fat bodies of these genotypes were dissected and lipidomic analyses were conducted. As expected, lower levels of TGs in *Axn*-depleted adipocytes were rescued by ectopic expression of *Lsd-1* and *Lsd-2* (Fig. 4m). In addition, fatty acyl chains in TGs, which represents the amount of each fatty acyl chain in the total TG pool, were reduced by Wnt signaling, while ectopic expression of *Lsd-1* and *Lsd-2* significantly rescued these defects (Fig. 4n). However, the level of FFAs in *Axn^RNAi^* adipocytes was reduced in this analysis (Fig. 4o). These results differed from analyses using the whole larvae (Fig. 2a) (Zhang et al., 2017), likely indicating that the bulk of FFAs were released from the larval adipocytes into the larval body cavity, thus not accounted for in the lipidomic analyses using the dissected fat body. Detailed changes in different types of TGs, fatty acyl chains in TGs, and FFAs are shown in Suppl Fig. S5. Together, these data show that increasing LDAPs, such as Lsd1 and Lsd2, rescues fat accumulation defects caused by active Wg signaling. This evidence suggests that Wg signaling can stimulate lipolysis and lipid mobilization by repressing the expression of LDAPs.

## Wg signaling directly represses the transcription of genes encoding the key factors required for the maintenance of intracellular lipid homeostasis

Active Wnt/Wg signaling is best known for stimulating the transcription of its target genes (Bejsovec, 2018; Nusse and Clevers, 2017). We investigated whether Wg signaling directly or indirectly regulates the transcription of these lipid metabolism-related genes. Specifically, we sought to identify the direct binding sites of Pan, the *Drosophila* TCF/LEF ortholog and the key transcriptional factor downstream of the canonical Wg signaling pathway (Bejsovec, 2018; Franz et al., 2017; Nusse and Clevers, 2017), in larval adipocytes. A lack of ChIP-grade antibodies for the key components of the Wg signaling pathway has been a major bottleneck in the investigation of the transcriptional regulation of Wg signaling (Ramakrishnan et al., 2018). To overcome this obstacle, we tagged the endogenous *pan* locus with EGFP using CRISPR-Cas9 gene editing. This *pan^EGFP^* line was validated by sequencing the genomic DNA, and the nuclear location of the Pan^EGFP^ protein was observed in different larval tissues (Suppl Fig. S6). The *pan^EGFP^* homozygotes are fully viable and fertile, demonstrating that the EGFP tag does not interfere with normal functions of Pan.

To map the genome-wide Pan-binding sites, we dissected wing discs from the third instar *pan^EGFP^* homozygous larvae and performed the CUT&RUN (Cleavage Under Targets & Release Using Nuclease) assay. This technique allows for high efficiency and reproducibility in obtaining high quality ChIP-Seq data with fewer cells than traditional ChIP-Seq analyses (Meers et al., 2019; Skene et al., 2018; Skene and Henikoff, 2017). A total of 16726 Pan-binding peaks in 4220 specific genetic loci were identified (Fig. 5a). Comparison of this list of genes with the genes identified from the RNA-Seq analyses using the *Axn^RNAi^* or *slmb^RNAi^* adipocytes led to the identification of 749 upregulated and 759 downregulated genes harboring Pan-binding sites (Fig. 5a), suggesting that these genes may be directly activated or repressed by Wg signaling. As expected, Pan binding sites were identified in known Wg target genes such as *nkd* (Fig. 5b), as well as *cycD*, *fz1*, *fz3*, *fz4*, *Notumn*, and *Wnt4* (Suppl Fig. S7).

**Fig. 5.**
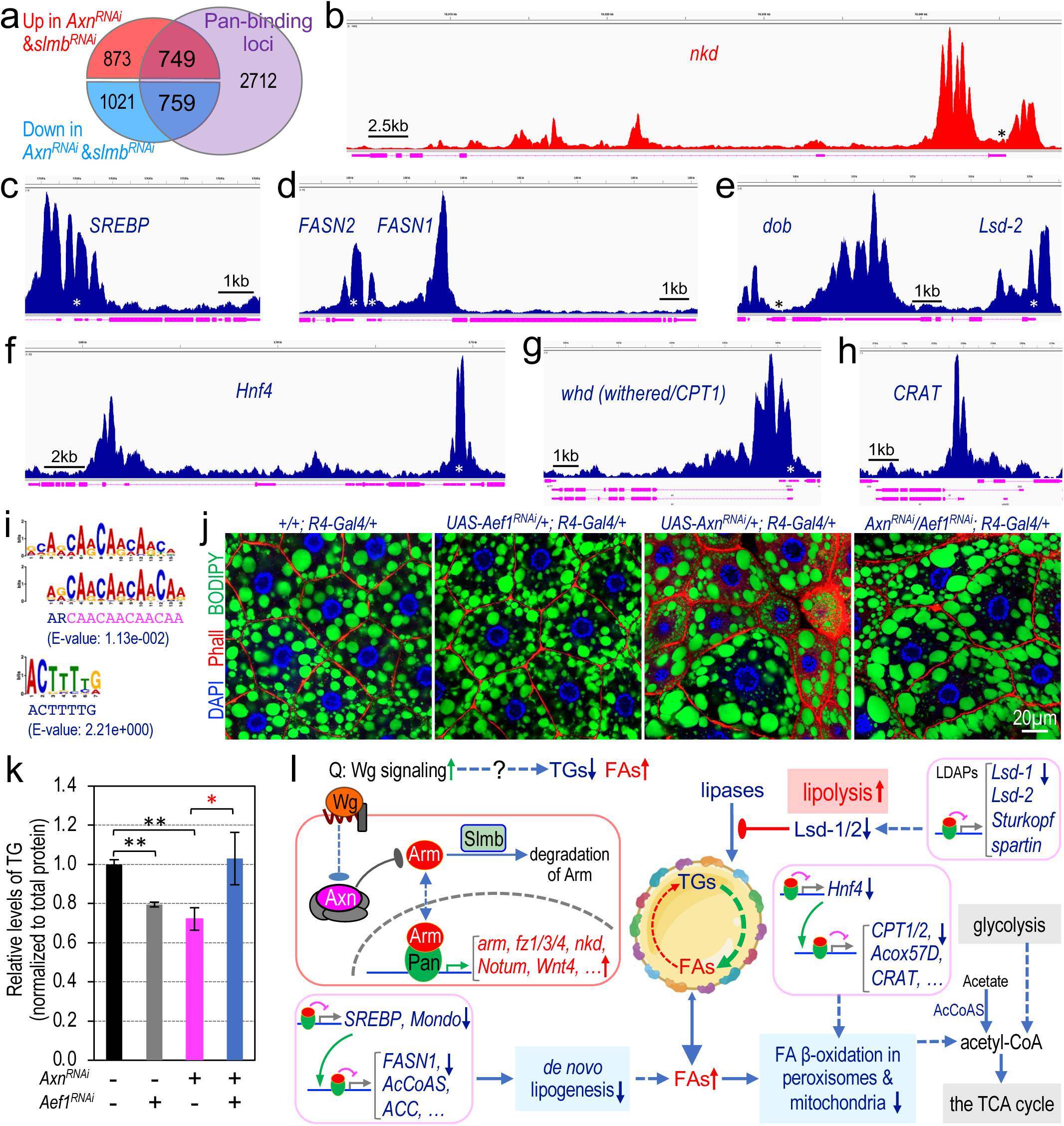
Wg signaling directly represses the transcription of key genes encoding factors required for lipogenesis, lipolysis, and LDAPs. (a) Venn diagram showing the number of genes altered in *Axn^RNAi^* or *slmb^RNAi^* fat body, as compared to the number of loci bound by Pan. (b-h) Genomic tracks are shown for Pan binding peaks at the following loci: *nkd* (b), *SREBP* (c), *FASN1* (d), *Lsd-2* (e), *Hnf4* (f), *whd* (*withered/CPT1*, g), and *CRAT* (h). The Reseq gene tracks are visualized using the Integrative Genomics Viewer (IGV) browser. The y-axis is autoscaled; the different isoforms of those genes are collapsed and shown in magenta; genes activated by Wg signaling are displayed in red track color, while genes downregulated by Wg signaling are shown in blue track color. The transcription start site (TSS) is indicated with an asterisk (*). (i) Enriched Pan-binding STREME motifs identified using the XSTREME tool and the STREME algorithms for motif discovery. (j) Co-depletion of *Aef1* rescues the fat accumulation defected caused by depleting *Axn* in larval adipocytes. Representative confocal images of larval adipocytes of the indicated genotypes stained with DAPI, Phall, and BODIPY. Scale bar: 20 μm. (k) Quantification of TG levels, and the results are normalized to the control (n = 3, independent biological repeats; * p<0.05; ** p<0.01). (l) A model depicting Wnt/Wg-driven lipid mobilization through repression of target gene transcription. Active Wnt signaling represses the transcription of Wnt/Wg target genes encoding key factors that regulate lipid mobilization such as Lsd-1 and Acox57D-d and lipogenesis such as FASN1. Lipid droplet-associated proteins (e.g., Lsd-1) hinder the access of lipases to triglycerides (TGs), diglycerides (DGs) and mono-glycerides (MGs). The colored dots on the surface of lipid droplets represent LDAPs. AcCoAS: Acetyl CoA synthase; Acox57D: acyl-CoA oxidase at 57D; Lsd-1: Lipid storage droplet-1.

Importantly, multiple Pan-binding peaks were revealed in genes encoding key lipogenic factors and enzymes such as *SREBP* and its target genes including *FASN1* (Fig. 5c-d), and other proteins involved in FA biosynthesis, such as *Mondo* (ChREBP homolog in flies), *AcCoAS*, *ACC* and others (Suppl Fig. S8a-h). As the mRNA levels of these genes were significantly reduced in the *Axn^RNAi^* and *slmb^RNAi^* adipocytes (Fig. 3g-3i), identification of Pan binding in the promoter regions of these key lipogenic genes suggests that Wg signaling inhibits lipogenesis by directly repressing the transcription of these lipogenic genes.

The second class of genes with direct Pan binding includes several LDAPs, such as *Lsd2* (Fig 5e), *Lsd2*, *spartin*, *sturkopf* and others (Suppl Fig. S8i-n). The third class of Wnt targets includes *Hnf4*, *CPT1* (*withered*/*whd*), and *CRAT* (Fig. 5f-5h), as well as *CPT2*, *ScpX* (sterol carrier protein X-related thiolase), *Acox57D* and other enzymes involved in FAO and electron transport chain (Suppl Fig. S9a-k). CPT1, CPT2, and CRAT are the transporters for importing FAs into the mitochondria and peroxisomes (Houten et al., 2020). Moreover, Pan also binds to the promoters of multiple genes encoding peroxisomal proteins, such as *Pex1* (encoding Peroxin 1), *Pex6*, *Pex13* and *ACOX1* (encoding acyl-CoA oxidase) (Suppl Fig. S9l-s). Therefore, increased lipolysis, together with reduced lipogenesis and FAO, constitutes a simple explanation for the general reduction of TGs and the increase of FFAs in adipocytes with active Wg signalin (Fig. 2) (Zhang et al., 2017).

## Wg signaling-induced repression of target gene transcription

Analyses of gene transcription driven by Wg signaling in different experimental systems often focus on target genes that are stimulated by Wg signaling, thus an unexpected finding of this study is the identification of a battery of Wg signaling repressed target genes in larval adipocytes. In contrast to three general principles underlying signaling-driven transactivation, e.g., activator insufficiency, cooperative activation, and default repression (Barolo and Posakony, 2002), “signal-induced repression” is an often-neglected general feature of the major signaling pathways in metazoans. This includes the Wg signaling pathway (Affolter et al., 2008). About 15-20 Wg signaling-repressed target genes have been characterized in *Drosophila* embryos, cultured Kc cells, and mice (Blauwkamp et al., 2008; Franz et al., 2017; Jamora et al., 2003; Piepenburg et al., 2000; Theisen et al., 2007). To compare the Pan binding sites for Wg-activated genes to Wg-repressed genes, we aligned normalized counts from 82 upregulated genes and 141 downregulated genes that have at least one enriched Pan-binding peak within their gene span. This analysis revealed that Pan is enriched around the transcription start sites (TSS), but no obvious difference was found in Wg-activated versus Wg-repressed target genes (Suppl. Fig. S10a).

Exactly how Wg signaling represses the transcription of these target genes remains poorly understood (Affolter et al., 2008; Hoverter and Waterman, 2008; Ramakrishnan et al., 2018). Analyses of the Pan binding sequence in the *Ugt36Bc* locus suggest that Wg repressed targets share the AGAWAW (W=A/T) consensus sequence for Pan binding, while Wg-activated targets contain the SCTTTGWW (S=G/C) motif. These Pan-binding motifs were proposed to allosterically define whether the Pan/Arm complex activates or represses target gene transcription (Blauwkamp et al., 2008). It is unclear if this motif represents a general feature of Wg target genes that are repressed upon Wg activation, because this sequence can be found frequently throughout the *Drosophila* genome (Blauwkamp et al., 2008). This problem has been difficult to address due to the lack of ChIP-grade antibodies (Ramakrishnan et al., 2018). Interestingly, our analyses of Pan-binding motifs of Wg repressed targets revealed two similar motifs that share the ARCAACAACAACAA (R=G/A) motif in 52% of the 759 genes (Fig. 5i). CAACAA motif is bound by a fat body-specific transcriptional repressor called Aef1 (Adult enhancer factor 1, a C2H2 Zinc finger protein) (Brodu et al., 2001; Falb and Maniatis, 1992a, b). Thus, we surmise that Aef1 may function in Wnt signaling-induced transcription repression of genes that regulate lipid homeostasis in fat body. A key prediction of this model is that depleting Aef1 in Wnt active adipocytes would alleviate the transcriptional repression of lipid catabolism-related genes by active Wnt signaling, thus should rescue the defects in fat accumulation. Indeed, co-depleting *Aef1* and *Axn* in larval adipocytes indeed strongly rescued the fat defects caused by depleting *Axn* alone (Fig. 5j), accompanied by the normalized levels of TG (Fig. 5k) and transcription of genes such as *ACC*, *FASN1*, and *Lsd-2* (Suppl. Fig. S10b-e). These results suggest that Aef1 may play a key role in mediating active Wnt signaling-induced transcriptional repression.

## Discussion

This study reveals that activated Wnt signaling reduces the levels of TGs in lipid droplets by inhibiting *de novo* lipogenesis, while promoting lipolysis in *Drosophila* larval adipocytes. Mechanistically, active Wg signaling attenuates lipogenesis by directly repressing the transcription of the key lipogenic enzymes, such as *FASN1*, *AcCoAS,* and *ACC.* Wg signaling may also repress the expression of *SREBP* and *Mondo* (ChREBP), thereby indirectly reducing the transcription of these key lipogenic enzymes (Fig. 5l). In parallel, Wg signaling also directly represses the transcription of genes encoding many LDAPs, such as *Lsd-1* and *Lsd-2* (Fig. 2j, 3d-f, 5e). Reduced LDAPs may facilitate lipases to access and convert TGs into FAs in LDs, thereby promoting lipid catabolism (Fig. 5l). Moreover, active Wg signaling directly inhibits the transcription of *Hnf4* and genes encoding key enzymes involved in FAO, such as *CPT1*, *CPT2*, *CRAT*, *Acox57D*, and *yip2*, thereby attenuating FAO in mitochondria and peroxisomes (Fig. 2h, Fig. 3, 5f-h, Suppl Fig. S9). Of these three processes, reduced *de novo* lipogenesis and increased lipolysis significantly reduces the accumulation of TGs in LDs, while increased lipolysis and simultaneous reduction of FAO increase the accumulation of FFAs in adipocytes. Synergistic effects among these three-pronged processes may reduce fat accumulation in Wg signaling-active adipocytes (Fig. 5l).

One unexpected finding of this study is that active Wg signaling directly inhibits the transcription of the key transcription factors and enzymes involved in lipogenesis, lipolysis, and FAO. The mechanism of this Wg signaling-induced transcriptional repression, instead of transcriptional activation, has been poorly understood. Our results suggest that the transcriptional repressor Aef1 may play an important role in Wg signaling-induced transcriptional repression, thereby controlling lipid homeostasis in adipocytes. Detailed molecular mechanisms warrants future investigations.

Our data show that the net effect of activated Wnt signaling is reduced accumulation of TGs and increased FFAs. Interestingly, it was recently reported that mammary tumor cells stimulated their adjacent adipocytes to convert TGs into FAs (Zhu et al., 2022). FAs can be absorbed by cancer cells for the biogenesis of phospholipids, the major membrane materials required for proliferation of cancer cells. Given that some of the Wnt targets include secreted Wnt ligands and Wnt signaling is aberrantly activated in nearly half of breast cancer cells, we speculate that cancer cells may fuel their proliferation by hijacking their neighboring adipocytes for lipid mobilization and releasing fatty acids through a Wnt signaling-dependent mechanism as reported in this study. If so, targeting enzymes required in lipid catabolism, combined with other chemotherapy or immunotherapy approaches, may offer synergistic advantages for cancer treatment.

## Methods

### Fly stocks and maintenance

Flies were maintained at 25°C on a standard cornmeal, molasses, and yeast medium. Only female larvae at the third instar wanderings stage were analyzed in this work. Information for the specific strains and their genotypes is included in Suppl Table 1.

### Cell biological analyses and Immunoblotting

Methods for larval adipocyte dissection, fixation conditions for immunostaining or staining with BODIPY, Nile Red, DAPI, and phalloidin were described previously (Zhang et al., 2017). Quantification of TG levels using thin layer chromatography (TLC) and a TG quantification colorimetric kit were performed as described previously (Li et al., 2022; Zhang et al., 2017).

For immunoblotting, cultured S2R+ cells were rinsed and resuspended in ice-cold 1xPBS then lysed in Tropix Lysis Solution (Applied Biosystems #2005117) with a protease inhibitor cocktail (Roche #11836170001), incubated on ice for 15 minutes. Lysates were centrifuged at 16,000Xg for 30 min at 4°C then protein concentrations were determined by Quick Start™ Bradford 1x Dye Reagent (Bio-Rad #5000205). For each sample, 20 μ loaded into 10% Mini-PROTEAN TGX Gels (Bio-Rad #4561034) in SDS running buffer then blotted onto nitrocellulose membranes through Trans-Blot Turbo RTA Mini 0.2 µm Nitrocellulose Transfer Kit (Bio-Rad #1704270). Membranes were hybridized with 1:1000 dilution of the anti-Arm primary antibody, purchased from the Developmental Studies Hybridoma Bank (DSHB Hybridoma Product N2 7A1 Armadillo, deposited by Eric Wieschaus), and 1:2000 of anti-Actin (Invitrogen #MA5-11869). 1:5000 dilution of anti-mouse-HRP-conjugated (Jackson ImmunoResearch) as the secondary antibody. Western Lightning Plus-ECL (Perkin Elmer LLC #NEL105001EA) was used for visualization.

### The HCR RNA-FISH assay

The methods for the multiplexed in situ HCR were described in detail previously (Li et al., 2022). The following probe sets and amplifiers were purchased from Molecular Instruments: the B1-Alexa Fluor 488 amplifiers were used with the probe sets for *FASN1* (lot number PRQ415), *Lsd-1* (PRQ418), and *arm* (PRR569); the B2-Alexa Fluor 594 amplifiers were used with the probe sets for *AcCoAS* (lot number PRQ416), *Lsd-2* (PRQ419), and *SREBP* (PRR570); and the B3-Alexa Fluor 647 amplifiers were used with the probe sets for *ACC* (lot number PRQ417), *Acox57D-d* (PRQ420), and Hnf4 (PRR571). Confocal images were taken using a Zeiss LSM900 confocal microscope system and then processed using Adobe Photoshop 2021.

### Cell culture and dual isotope radiolabeling

*Drosophila* S2R+ cells (DGRC stock #150; https://dgrc.bio.indiana.edu//stock/150 ; RRID:CVCL_Z831) and S2-Tub-wg (DGRC stock #165; https://dgrc.bio.indiana.edu//stock/165 ; RRID:CVCL_1B57) were purchased from the *Drosophila* Genomics Resource Center (DGRC) and cultured in Schneider’s *Drosophila* Medium (Gibco #21720-024) supplemented with 10% FBS (ATCC #30-2020) and incubated at 25°C. Wg-conditioned medium (WCM) and Control media (CM) were harvested from S2-Tub-wg and S2R+ cells respectively, which were seeded 24 h prior to collecting the supernatant (1x10^6^ cells/ml for immunoblotting and 5×10^6^ cells/ml for dual isotope radiolabeling experiments described below) by centrifuging the cells at 500xg for 5 minutes and filtered with 0.45 μm sterile syringe filters (Millex-HV #SLGV033RS). For dual isotope radiolabeling experiments, S2R+ cells were seeded in six-well plates (2 ml/well; 3.5×10^6^ cells/ml) for 24h together with radioactively labeled [^14^C(U)]D-glucose (5 μl/well; 6.0 10^8^ dpm/μmol) and [9,10- ^3^H(N)] Palmitic acid (2 μl/well; 1.05 10^11^ dpm/μmol). Next, the experimental groups and control groups were treated with Wg-conditioned medium (WCM) or control media (CM) respectively. WCM was harvested from S2 tubulin wingless cells which were seeded 24 h prior to collecting the supernatant (5×10^6^ cells/ml) by centrifuging the cells at 3500 rpm for 5 min. For the control medium, S2R+ cells were prepared as described above. S2R+ cells were collected from CM and WCM treated wells respectively at 0hr, 12hr and 24hr. Collected cell pellets were washed with (1x) PBS and TLC was performed as previously described (Hildebrandt et al., 2011; Zhang et al., 2017), with the following minor modifications. The specific regions corresponding to FFAs and TGs were scrapped out of the TLC plates and collected into scintillation vials. ^3^H and ^14^C cpm (counts per minute) values were determined by counting in 4 ml of CytoScint™ Liquid Scintillation Cocktail (Catalog number: 882453) in a Hidex 300 SL™ automatic liquid scintillation counter. The dpm (disintegrations per minute) values for ^3^H and ^14^C were computed by using NIST (National Institute of Standards and Technology) traceable standards. The efficiencies of ^3^H and ^14^C counting were 33.06% and 87.23% respectively for an optimized period of 100 seconds measuring time. Dpm values for ^14^C and ^3^H within TG and FFA were calculated using the respective efficiency figures. ^14^C/^3^H ratios were calculated and, normalized ratios were plotted.

### RNA-seq sample preparation, sequencing and gene ontology enrichment analysis

The total RNA from dissected fat bodies of 20 third-instar larvae was extracted for each biological replicate using 1.0 ml of TRIzol Regent (Invitrogen), and then purified using the RNeasy Mini Kit (Qiagen, 74104) following the manufacturer’s instructions. The ribosome depletion stranded RNA library was generated by using TruSeq Stranded Total RNA Library Prep kit (Illumina, 20020596) according to the manufacturer’s manual, and the RNA libraries were sequenced by the BGI Americas Corporation. The sequences were aligned to the Dm6 reference genome by RNASTAR (Widmann et al., 2012). The count table was generated with featureCounts from Subread (Liao et al., 2014). The differential gene analysis within each group of experiments was performed by DESeq2 and the normalized count table was generated by DESeq2 (Love et al., 2014). Differentially expressed genes had adjusted p-value < 0.05. ClusterProfiler and DAVID GO were used for gene ontology enrichment analyses (Sherman et al., 2022; Wu et al., 2021), and the results were visualized using ggplot2 (Wickham, 2016). The heatmaps were generated by complexHeatmap (Gu, 2022).

### Quantitative proteomics analysis

Protein digestion was performed using the filter-aided proteome preparation (FASP) method with slight modifications (Wisniewski et al., 2009). Briefly, the proteins were reduced with 100 mM DTT at 37 °C for 1 h, and the lysates were transferred to the Microcon YM-30 centrifugal filter units (EMD Millipore Corporation, Billerica, MA). The lysis buffer was replaced with 200ul UA (8M Urea, 100mM Tris.Cl pH8.5) twice. After being alkylated by 55 mM iodoacetamide (IAA, Sigma-Aldrich, Saint Louis, MO), the denaturing buffer was replaced by a buffer containing 0.1 M triethylammonium bicarbonate (TEAB, Sigma-Aldrich, Saint Louis, MO). Proteins were then digested with sequencing grade trypsin (Promega, Madison, WI) at 37 °C overnight, and the resultant tryptic peptides were labeled with acetonitrile-dissolved TMT reagents (Thermo Scientific, Rockford, IL), and equal amounts of labeled samples were mixed before prefractionation with reversed phase (RP)-high performance liquid chromatography (HPLC). Briefly, the peptides were fractionated on a phenomenex gemini-NX 5u C18 column (250 x 3.0 mm, 110 Å) (Torrance, CA, USA) using a Waters e2695 separations HPLC system. The separated samples were collected and combined into 10 fractions. All fractions were dried with a Speed-Vac concentrator and desalted by StageTip (Rappsilber et al., 2007). The LC-MS/MS analysis was performed using an LTQ Orbitrap Elite mass spectrometer (Thermo Scientific, Rockford, IL Waltham, MA) coupled online to an Easy-nLC 1000 in the data-dependent mode. The peptides were separated on a capillary analytic column (length: 25 cm, inner diameter: 75 μm (diameter: 5m μ). The positive ion mode was used for MS measurements, and the spectra were acquired across the mass range of 300-1800 m/z. Higher-energy collisional dissociation (HCD) was used to fragment the fifteen most intense ions from each MS scan. The database search was performed for all raw MS files using the software MaxQuant (version 1.5). The *Drosophila melanogaster* proteome sequence database downloaded from uniprot (https://www.uniprot.org/) was applied to search the data. The parameters used for the database search were set up as follows: The type of search: MS2; The protease used for protein digestion: trypsin; The type of isobaric labels: 6-plex TMT; The minimum score for unmodified peptides: 15. Default values were used for all other parameters.

### Lipid sample preparation and shotgun lipidomics analysis

Dissected larval fat bodies were snap frozen, then pulverized using a liquid nitrogen-cooled mortar and pestle. The pulverized samples were further homogenized in 10-times diluted PBS using a Precellys Evolution^®^ Homogenizer at 6000 rpm for 20 s and paused for 10 s at 4 °C for 3 consecutive cycles. Protein assay on individual homogenates was conducted. An aliquot of the homogenate was transferred to a disposable glass test tube. A mixture of lipid internal standards for quantification of all reported lipid classes was added to the tube based on the tissue protein content; lipid extraction was performed by a modified Bligh and Dyer method as previously described (Wang and Han, 2014). All the lipid extracts were flushed with N_2_, capped, and stored at -20 °C.

For ESI-MS analysis after direct infusion, individual lipid extracts were further diluted to a final concentration of ∼500 fmol/µL, and the mass spectrometric analysis was performed on a Q-Exacutive mass spectrometer (Thermo, San Jose, CA) for cardiolipin, lyso-cardiolipin, and PG analysis and on a QqQ mass spectrometer (Thermo TSQ Altis, San Jose, CA) for other lipids. Both of instruments were equipped with an automated nanospray device (TriVersa NanoMate, Advion Bioscience Ltd., Ithaca, NY) to ionize the lipid species by nano-ESI and operated with Xcalibur software as previously described (Yang et al., 2009). Identification and quantification of all lipid molecular species of interest were performed using an in-house automated software program following the principles for quantification by mass spectrometry as previously described in detail (Yang et al., 2009). Fatty acyl chains of lipids were identified and quantified by neutral loss scans or precursor ion scans of corresponding acyl chains and calculated using the same in-house software program (Yang et al., 2009). All lipid levels were normalized to sample protein content.

### CRISPR Cas9 mediated tagging of Pangolin with EGFP

Pangolin gene structure information was retrieved from FlyBase (FBgn0085432), and primary amino acid sequence was retrieved from Uniprot (Q8IMA8). After analyzing the domain structure of the protein (https://www.ebi.ac.uk/interpro/result/InterProScan/ iprscan5-R20220926-005559-0039-31654548-p1m/), we decided to introduce the EGFP tag at the C-terminus of Pangolin. Using CRISPR Optimal target finder (http://targetfinder.flycrispr.neuro.brown.edu), two sgRNAs were located within the last intron and 3’-UTR regions, respectively. Those two sgRNAs were cloned into the pCFD3 vector (http://www.crisprflydesign.org/plasmids/) (Port et al., 2014). Primer designing was performed by using NEB Builder (https://nebuilder.neb.com/#!/; primer sequences are listed in Table S2). NEBuilder HiFi DNA Assembly Cloning Kit (NEB #E5520) was used to introduce upstream and downstream homology arms (1000bp each) and the last intron-exon regions together with the EGFP coding region into a donor vector. The BAC clone ‘BACR21B19’ was obtained from the BACPAC Resources Center and used as template for PCR. The donor vector and two sgRNA constructs in PCDF3 were embryo injected. The genetic screening of the transgenic flies carrying endogenous EGFP tag was performed by using classical fly genetic approaches and PCR validations using extracted genomic DNA.

### CUT&RUN sequencing to identify the genome-wide Pan-binding sites

For each sample, 20 wing discs from the third instar wandering larvae were dissected in Schneider medium and then transferred into an Eppendorf tube containing 100 μL 1X Wash buffer (CUT&RUN assay kit #86652, CST). Next, 10 μL of activated Concanavalin A Magnetic bead suspension was added to the wing disc, and the tube rotated at room temperature for 10 min. The Eppendorf tubes were placed on the Magnetic rack and the beads were resuspended by adding 100 μL DBE buffer (from the kit) and rotated for 10 min at room temperature. 100 μL Antibody binding buffer containing the appropriate amount of primary antibody (anti-GFP, ab290), or negative control (rabbit mAb IgG from the kit), was added. The Eppendorf tubes were placed on a nutator overnight at 4 °C. The remaining steps were carried out by following the manufacturer’s instructions on the kit. The extracted DNA was purified by using DNA purification buffers and spin columns (#14209S, CST). The DNA libraries were constructed using the TruSeq ChIP SMP Prep kit (IP-202-1012, 15034288, 15027084, Illumina) by following the sample preparation guide. The sequencing data was processed with nf-core/cutandrun v2.0 (Cheshire and West), an integrated pipeline for the CUT&RUN assay. The marked alignments from the pipeline were further processed for peak calling and normalization with MACS3 (https://macs3-project.github.io/MACS/). The averaged and normalized pileup bedgraph file from MACS3 was converted to a bigwig file with bedGraphicToBigWig from UCSC-toolkit for visualization in IGV. The profile of Pan binding was plotted with deepTools. To discover the motifs in Pan-binding sites, we extracted the sequences +100bp of called summits within the genes of interest, and then used the XSTREME tool with default settings for *de novo* motif discovery (Grant, 2021).

### Statistical analyses

P-values were calculated based on one-tailed unpaired *t*-tests, and error bars indicate standard deviation. At least three independent biological repeats were analyzed for each experiment shown in Fig. 1i, 1k, 2a, 4g, 4m-o, and Suppl Fig. 6. * p<0.05; ** p<0.01; *** p<0.001.

### Data Availability

The RNA-seq data and sequencing results from the CUT&TUN assay will be deposited in NCBI’s Gene Expression Omnibus upon acceptance of this work.

## Supporting information

Table S1 List of Stocks and Primers

Table S2 Primers used

## Acknowledgements

We thank Yashi Ahmed and Mathias Beller for sharing fly strains, Kevin Charbonneau for his advice on using the liquid scintillator, and Fu-Ning Hsu for preparing samples used in the proteomic analyses. We thank the Bloomington *Drosophila* Stock Center (NIH Grant P40OD018537) for fly stocks, the *Drosophila* Genomics Resource Center (NIH Grant 2P40OD010949), and the Developmental Studies Hybridoma Bank at the University of Iowa for monoclonal antibodies. We are also grateful to Keith Maggert, Robert Schofner and Jasmine Sun for critical comments on the manuscript. The Functional Lipidomics Core at Barshop Institute for Longevity and Aging Studies is partially supported by the National Institute on Aging grants P30 AG013319 and P30 AG044271 (to X. Han). This research was supported by a grant from the National Institute of Health (GM129266 to J.-Y.J.).

## Author Contributions

Conceptualization was done by M.L., R.-U.-S. H.-W., and J.-Y.J. Methodology was developed by M.L., R.-U.-S. H.-W., and X.L. Investigation was done by M.L., R.-U.-S. H.-W., X.L., X. Huang, T.-H. L., and X. Han. Formal analysis was carried out by M.L., R.-U.-S. H.-W., X.L., X. Huang, X. Han, Y.W., and J.-Y.J. Data were curated by M.L., R.-U.-S. H.-W., X.L., and J.-Y.J. The original draft was written by M.L., R.-U.-S. H.-W., and J.-Y.J. Review and editing of the draft were done by M.L., R.-U.-S. H.-W., X. Han, Y.W., and J.-Y.J. Visualization was done by M.L., R.-U.-S. H.-W., and J.-Y.J. Supervision was the responsibility of J.-Y.J. Project administration was done by J.-Y.J. Funding was acquired by X. Han and J.-Y.J.

### Materials & Correspondence

**Correspondence and requests for materials should be addressed to J.-Y.J.**

**Suppl Fig. S1. Effects of Wg signaling on larval adipocytes.** (a-f) Depleting Axn or Slmb stabilizes Arm proteins. (a-c) Representative confocal images of larval adipocyte from larvae of indicated genotypes immunostained with Arm antibody (red) and DAPI (blue). Genotypes: (a) *SREBP-Gal4/+; +*; (b) *SREBP-Gal4/UAS-Axn^RNAi^; +*; and (c) *SREBP-Gal4/+; UAS-slmb^RNAi^/+*. (d-f) Confocal images of larval adipocytes from female larvae of indicated genotypes stained with DAPI (blue). Green: eGFP-tagged Arm. Genotypes: (d) *arm^EGFP^/+; SREBP-Gal4/+; +*; (e) *arm^EGFP^/+; SREBP-Gal4/UAS-Axn^RNAi^; +*; and (f) *arm^EGFP^/+; SREBP-Gal4/UAS-Axn^RNAi^; UAS-slmb^RNAi^/+*. Scale bar in c: 20 μm. (g-h) Depleting Axn stimulates the expression of Wg target gene *nkd* in larval adipocytes. Larval adipocytes were stained with Phall (red) and DAPI (blue). Genotypes: (g/g’) *+; R4-Gal4/nkd^EGFP^*; (h/h’) *UAS-Axn^RNAi^/+; R4-Gal4/nkd^EGFP^*. Note the elevated Nkd^EGFP^ in small adipocytes than the big adipocytes marked with asterisks (*). Scale bar in g’: 20 μm. (i-o) Active Wg signaling disrupts lipid accumulation in larval adipocytes. (i) Quantification of TG levels of the indicated genotypes (n = 3 biological repeats). Results are normalized to control (*dCg-Gal4/+; +*). (j-m) Depleting Axn using different fat body-specific Gal4 lines and two *UAS-Axn^RNAi^* lines (on the second and third chromosomes) led to similar defects in lipid accumulation. Genotypes: (j) *FB-Gal4/+; +*; (k) *FB-Gal4/UAS-Axn^RNAi^; +*; (l) *+; SREBP-Gal4/+*; and (m) *UAS-Axn^RNAi^/+; SREBP-Gal4/+*. (n-o) Knocking down Pan rescues the defects in lipid accumulation caused by depletion of Axn. Genotypes: (n) *UAS-pan^RNAi^/+; SREBP-Gal4/+*; and (o) *UAS-pan^RNAi^/UAS-Axn^RNAi^; SREBP-Gal4/+*. Scale bar in m: 20 μm.

**Suppl Fig. S2. Principle of the dual isotope radiolabeling experiment.** (a) Key metabolic processes of radiolabeled [^14^C(U)] D-glucose and [9,10-^3^H(N)] Palmitic acid. ^14^C is marked with red stars and ^3^H is highlighted in blue. (b) Separating radiolabeled TGs and FFAs using TLC. The levels of ^14^C and ^3^H in TGs and FFAs were measured using a liquid scintillation counter, and the ^14^C/^3^H ratios were calculated in TGs and FFAs.

**Suppl Fig. S3. Bioinformatic analyses of the RNA-seq data for genes that are altered in *Axn^RNAi^* or *slmb^RNAi^* fat bodies.** (a) Heatmap of gene expression for genes related to the ‘Wnt signaling’ pathway. (b,c) Pathway enrichment bubble plots showing the major pathways that are significantly altered in larval fat body with the depletion of either *Axn* (b) or *slmb* (c). Genotypes: (b) *SREBP-Gal4/UAS-Axn^RNAi^; +*; and (c) *SREBP-Gal4/+; UAS-slmb^RNAi^/+.* (d) Heatmap of gene expression for genes related to the ‘Biosynthesis of unsaturated fatty acids’ pathway, note that the expression of most genes in this pathway are downregulated in *Axn^RNAi^* or *slmb^RNAi^* fat body. (e) Heatmap of gene expression for genes related to the ‘Peroxisome’ pathway.

**Suppl Fig. S4.** Representative confocal images of larval adipocytes stained with DAPI (blue) and LipidTox (red). Genotypes: (a) *dCg-Gal4/+; UAS-Lsd1^eGFP^/+*; (b) *dCg-Gal4/UAS-Axn^RNAi^; UAS-Lsd1^eGFP^/+*; (c) *dCg-Gal4/+; UAS-Lsd2^eGFP^*; and (d) *dCg-Gal4/UAS-Axn^RNAi^; UAS-Lsd2^eGFP^/+*.

**Suppl Fig. S5. Additional results from quantitative lipidomics measurement of different types of lipids in fat body dissected from the third instar larvae.** (a-b) Triglycerides (TGs), (c) The amount of fatty acyl chains in total TG pool, and (d) FFAs. Specific genotypes are color coded and shown below (f). * or #: p<0.05; ** or ##: p<0.01; comparisons between ‘*dCg-Gal4/+; +*’ and ‘*dCg-Gal4/UAS-Axn^RNAi^; +*’ are indicated with asterisks (* or **), and comparisons between ‘*dCg-Gal4/+; UAS-Lsd1^eGFP^/UAS-Lsd2^eGFP^*’ and ‘*dCg/UAS-Axn^RNAi^; UAS-Lsd1^eGFP^/UAS-Lsd2^eGFP^*’ are indicated with # or ##.

**Suppl Fig. S6. Generation and validation of the *pan^EGFP^ Drosophila* strain.** (a) Design of the donor template using the pGEM-T-pan^EGFP^ vector. (b, c) Validation of the *pan^EGFP^* line using the genomic DNA from *pan^EGFP^* homozygous line by PCR (b) and sequencing (c). Pan is localized in the nuclei of larval fat body (d/d’) and wing imaginal discs (e/e’). (d’/e’) merged images with DAPI staining shown in blue. Scale bar in d’: 10 μ

**Suppl Fig. S7.** Tracks are shown for Pan binding at loci encoding factors related to the Wg signaling pathway. Genes activated by Wg signaling are displayed in red track color, and genes downregulated by Wg signaling are shown in blue track color. The transcription start site (TSS) is indicated with an asterisk (*). The y-axis is autoscaled; the different isoforms of those genes are collapsed and shown in magenta below each track.

**Suppl Fig. S8.** Tracks are shown for Pan binding at loci encoding factors related to the fatty acid biosynthesis pathway (a-h), and the loci encoding factors related to the LDAPs (i-n). Genes activated by Wg signaling are displayed in red track color, and genes downregulated by Wg signaling are shown in blue track color.

**Suppl Fig. S9.** Tracks are shown for Pan binding at loci encoding factors related to FAO and electron transport chain (a-k), as well as a few loci encoding factors related to peroxisome (l-s). The expression of these genes is downregulated by Wg signaling, and track color is shown in blue.

**Suppl Fig. S10. Analyses of the effects of Aef1 on gene expression.** (a) Alignment of normalized counts from 82 Pan-binding sites of upregulated genes and 141 Pan-binding sites of downregulated genes. Heatmap below shown the relative position of Pan-binding peaks within their gene spans. (b-e) The mRNA transcripts of *FASN1* (green), *ACC* (magenta) and *Lsd-2* (orange) in larval adipocytes detected by the HCR assay. Note the lower levels of *FASN1*, *ACC*, and *Lsd-2* in small adipocytes (indicated with asterisks) than the neighboring large adipocytes when Axn is depleted (d). These differences are rescued by co-depletion of *Aef1* (e). Genotypes: (b) +; *R4-Gal4/+*; (c) *UAS-Aef1^RNAi^/+; R4-Gal4/+*; (d) *UAS-Axn^RNAi^/+; R4-Gal4/+*; and (e) *UAS-Axn^RNAi^/UAS-Aef1^RNAi^; R4-Gal4/+*. Scale bar in d: 20 μm.

**Table S1 List of the *Drosophila* stocks used in this study.**

**Table S2 Primers used to generate the *pan^EGFP^* strain using the CRISPR-Cas9 technique.**

