## Supplementary figures and images for "Wingless signaling promotes lipid mobilization through signal-induced transcriptional repression"

### Suppl. Figures S1-S10

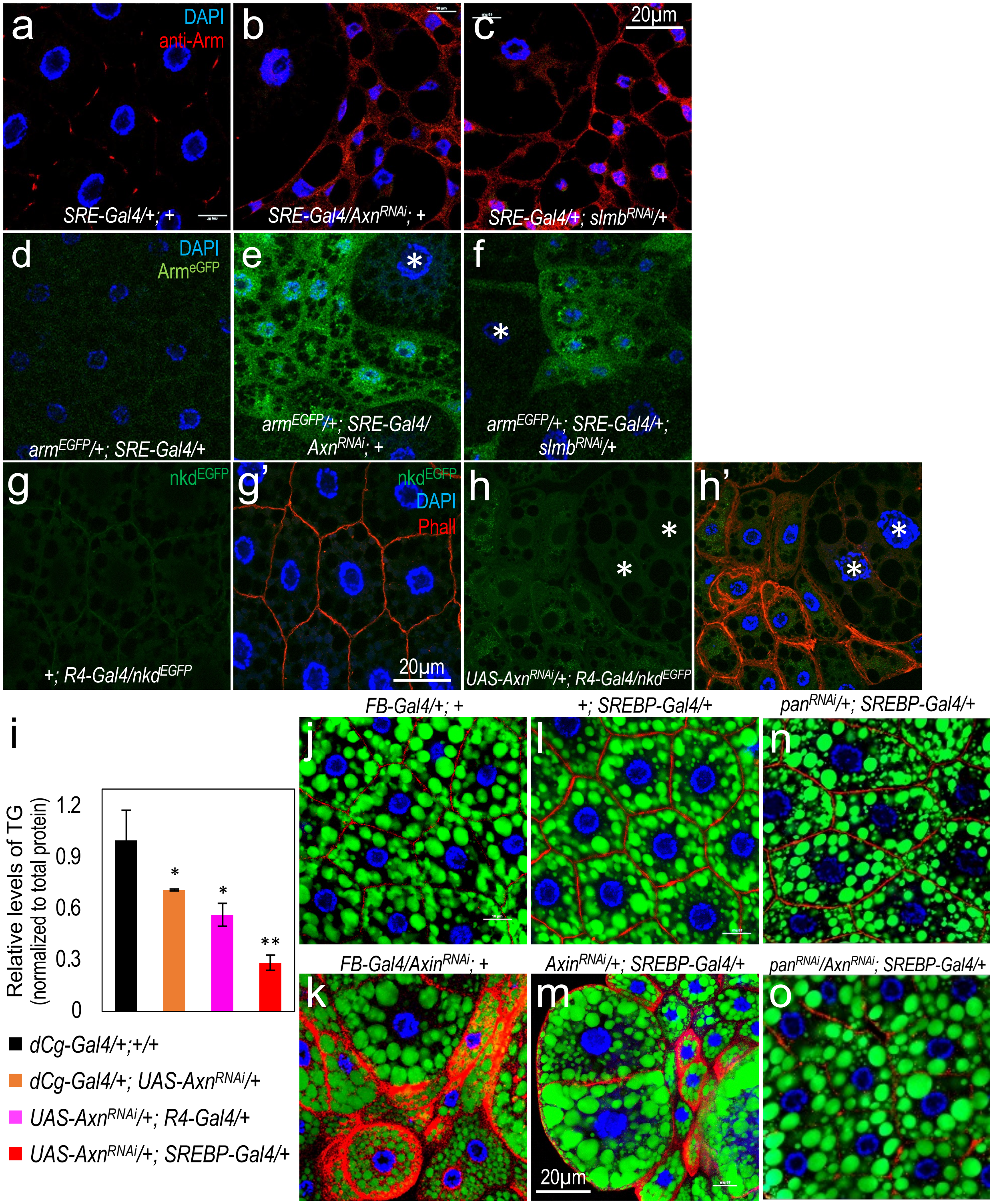

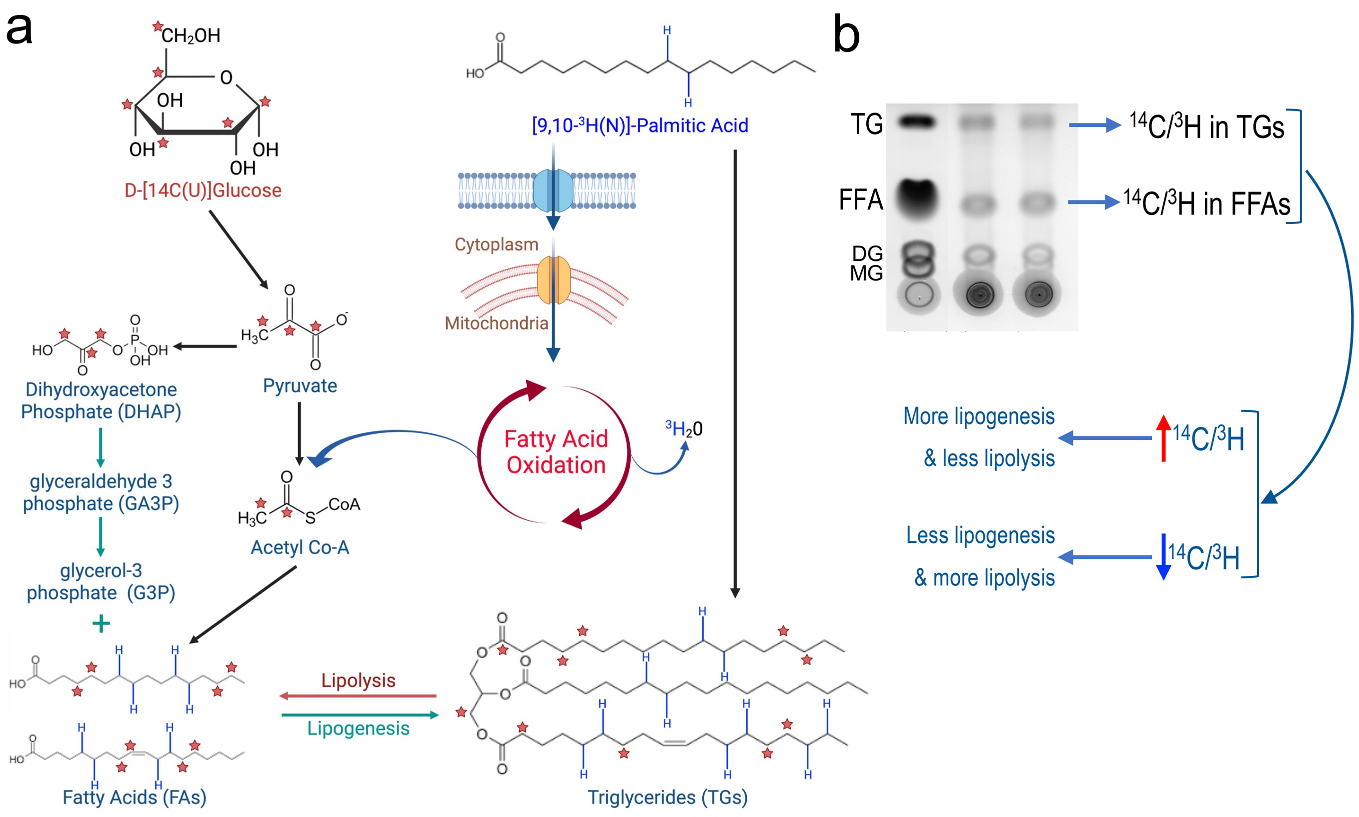

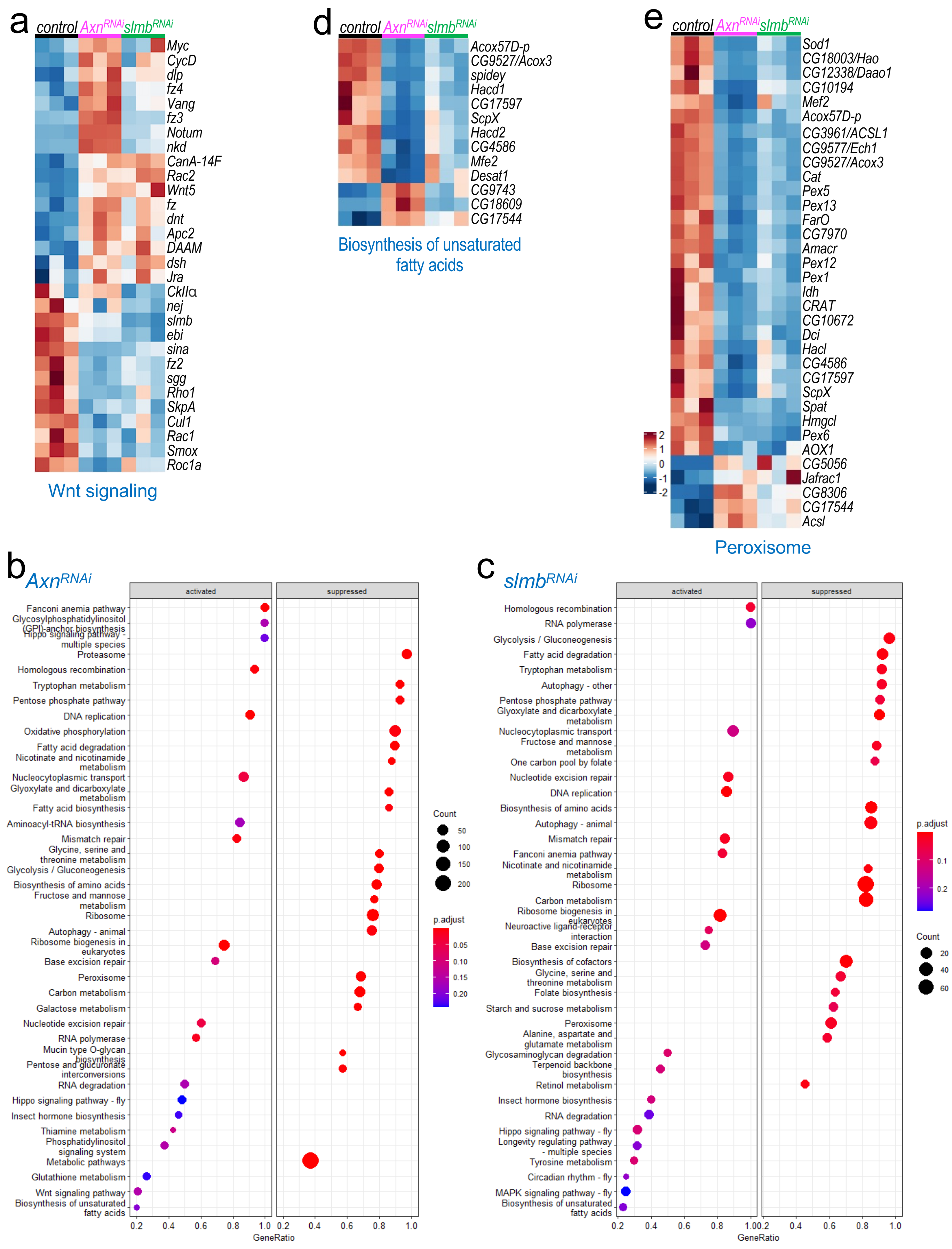

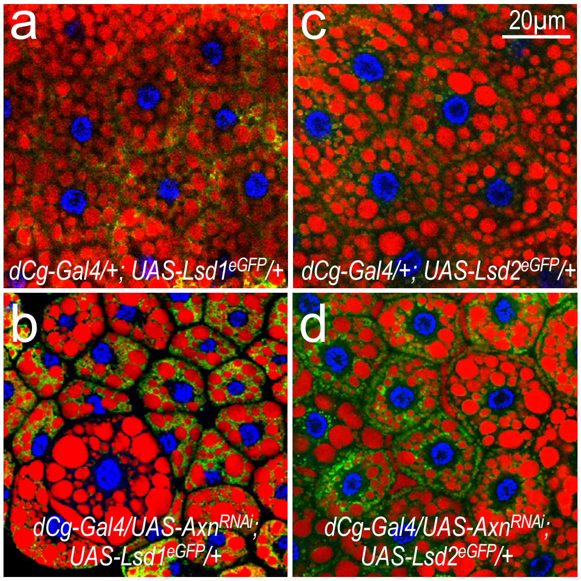

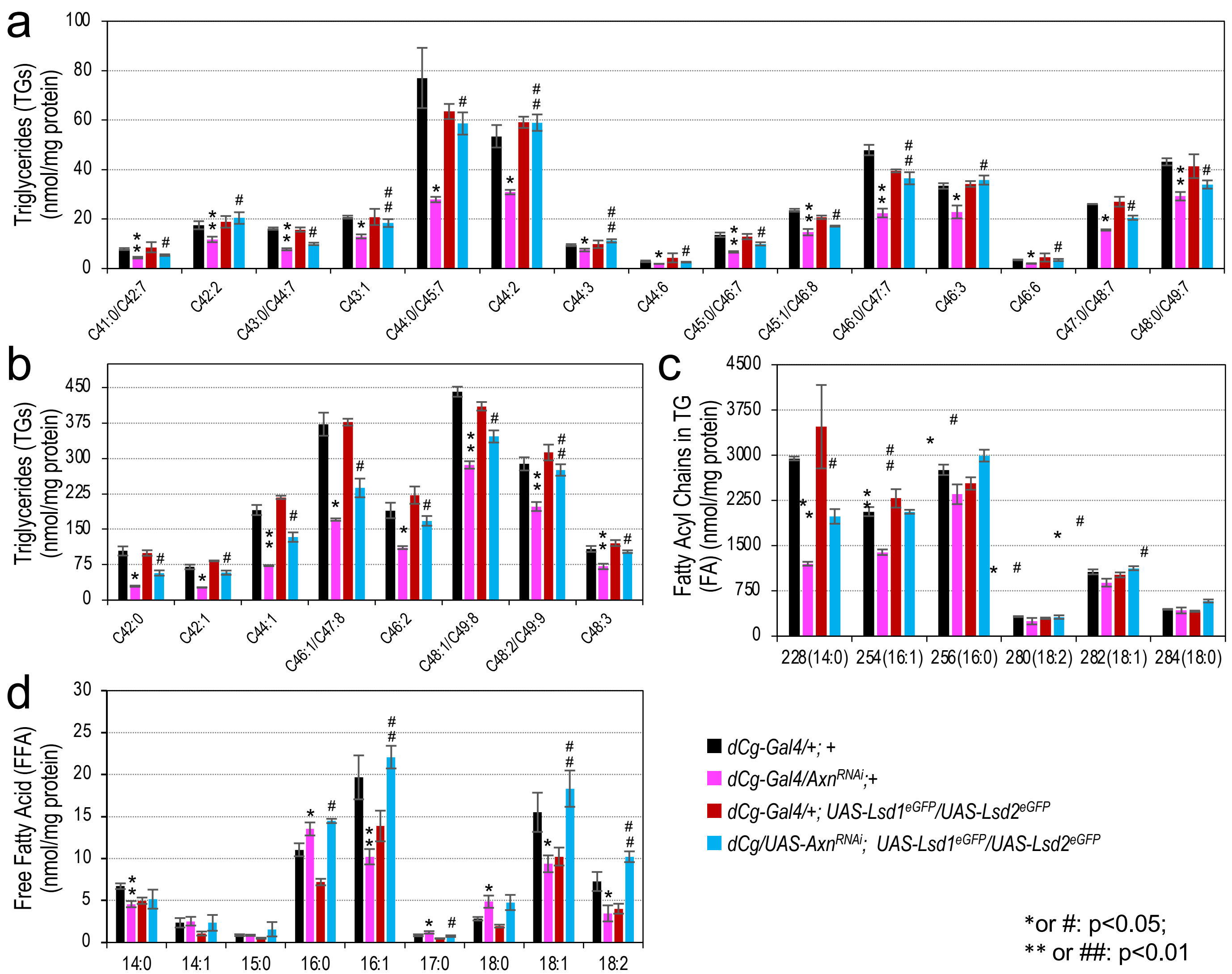

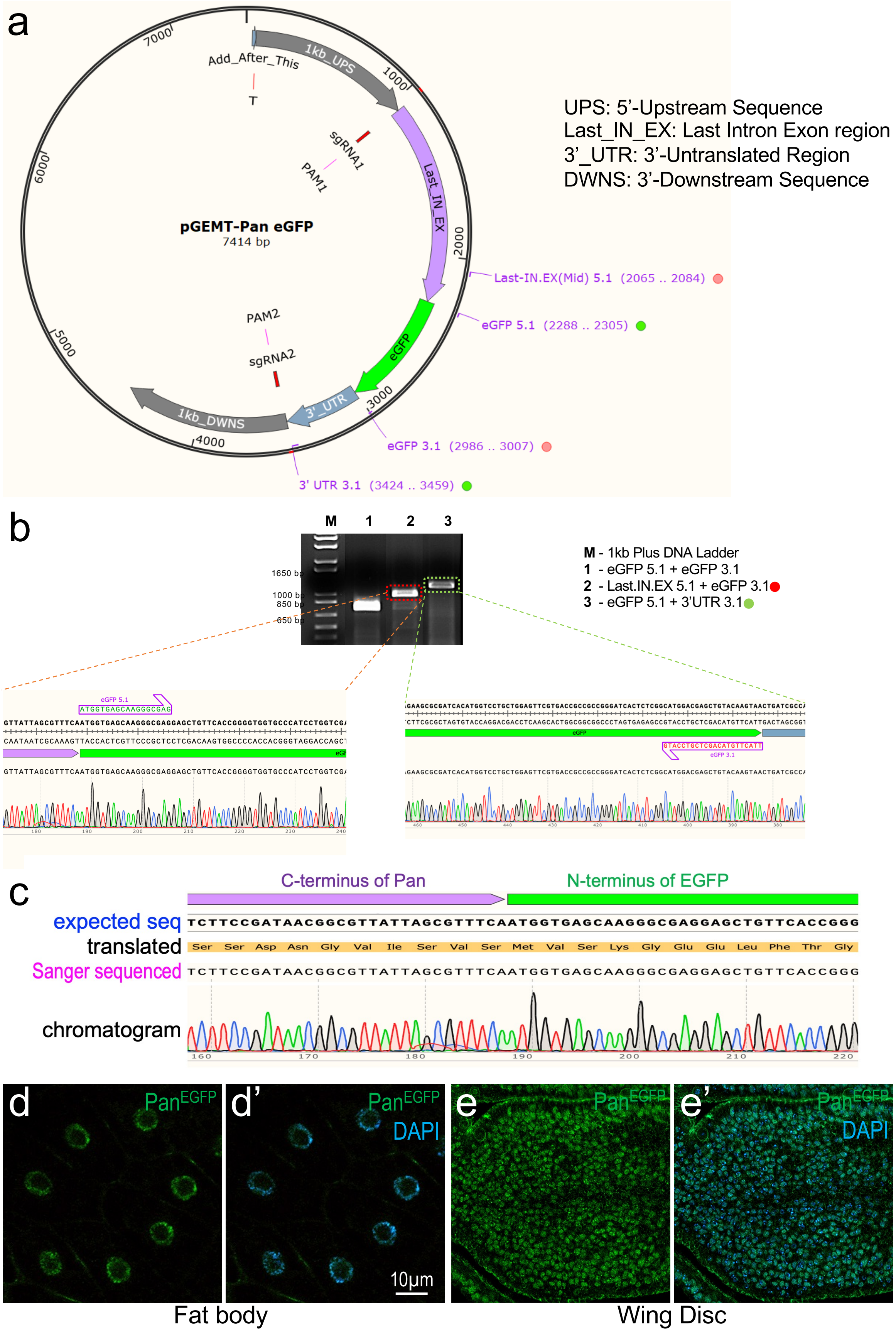

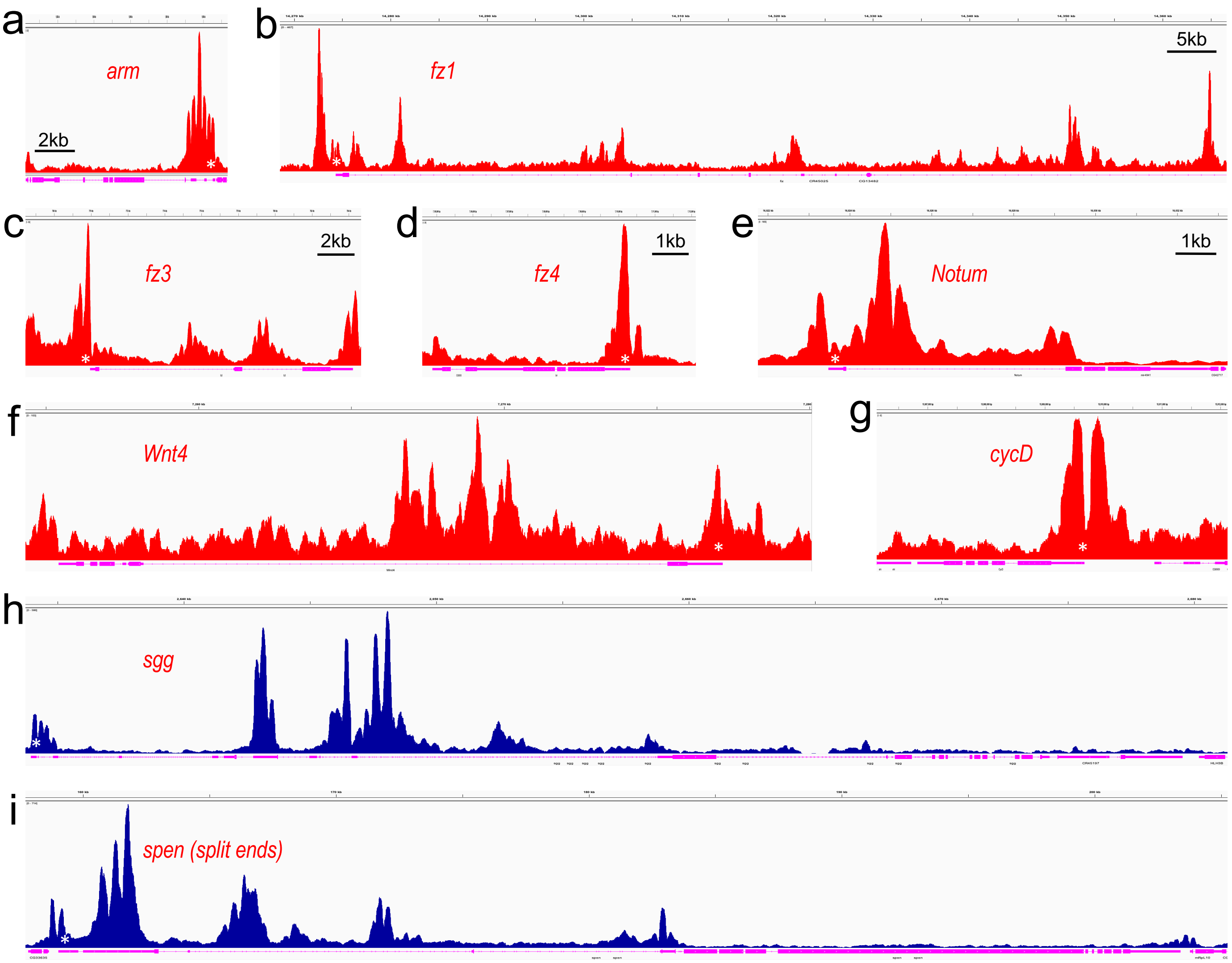

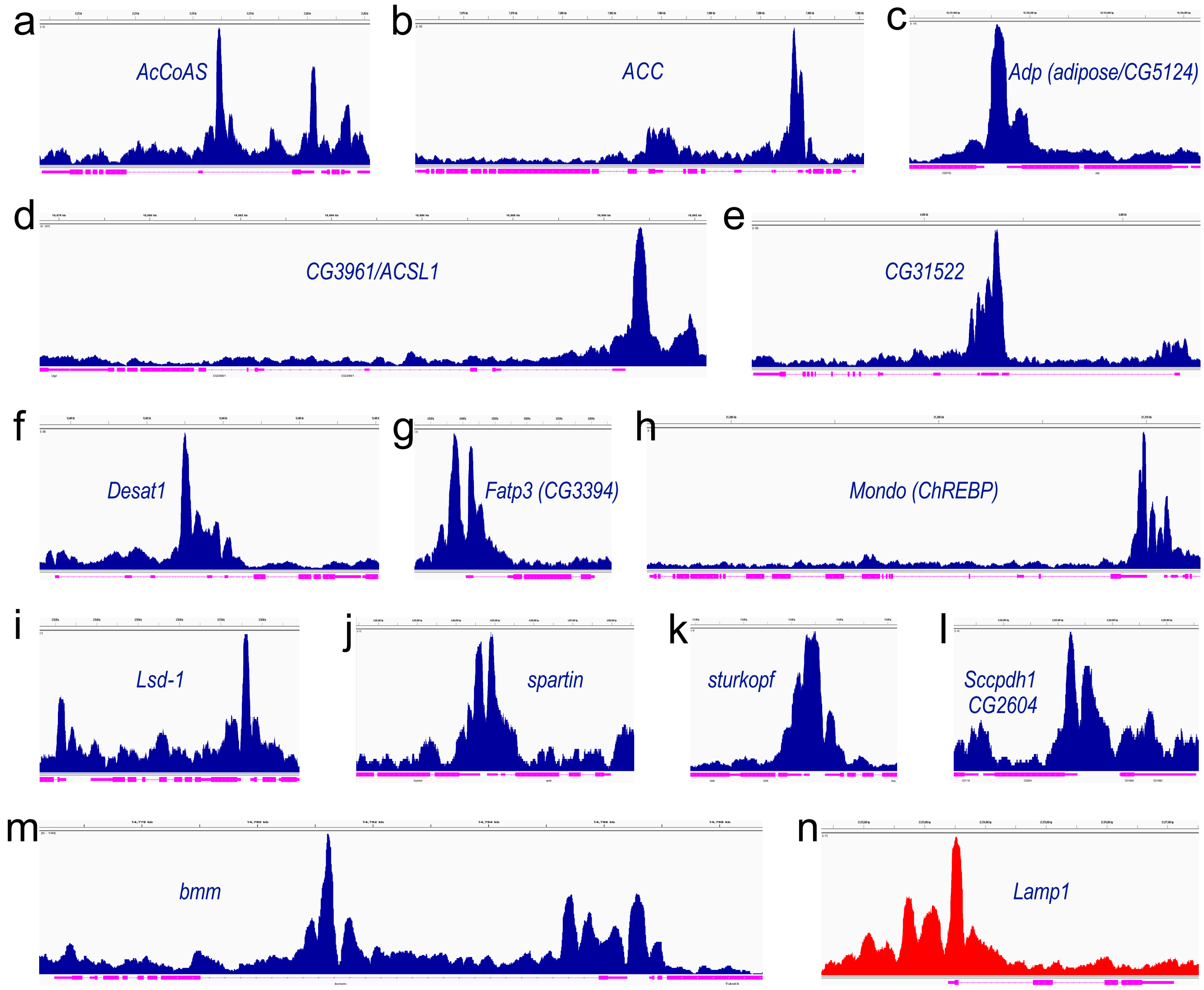

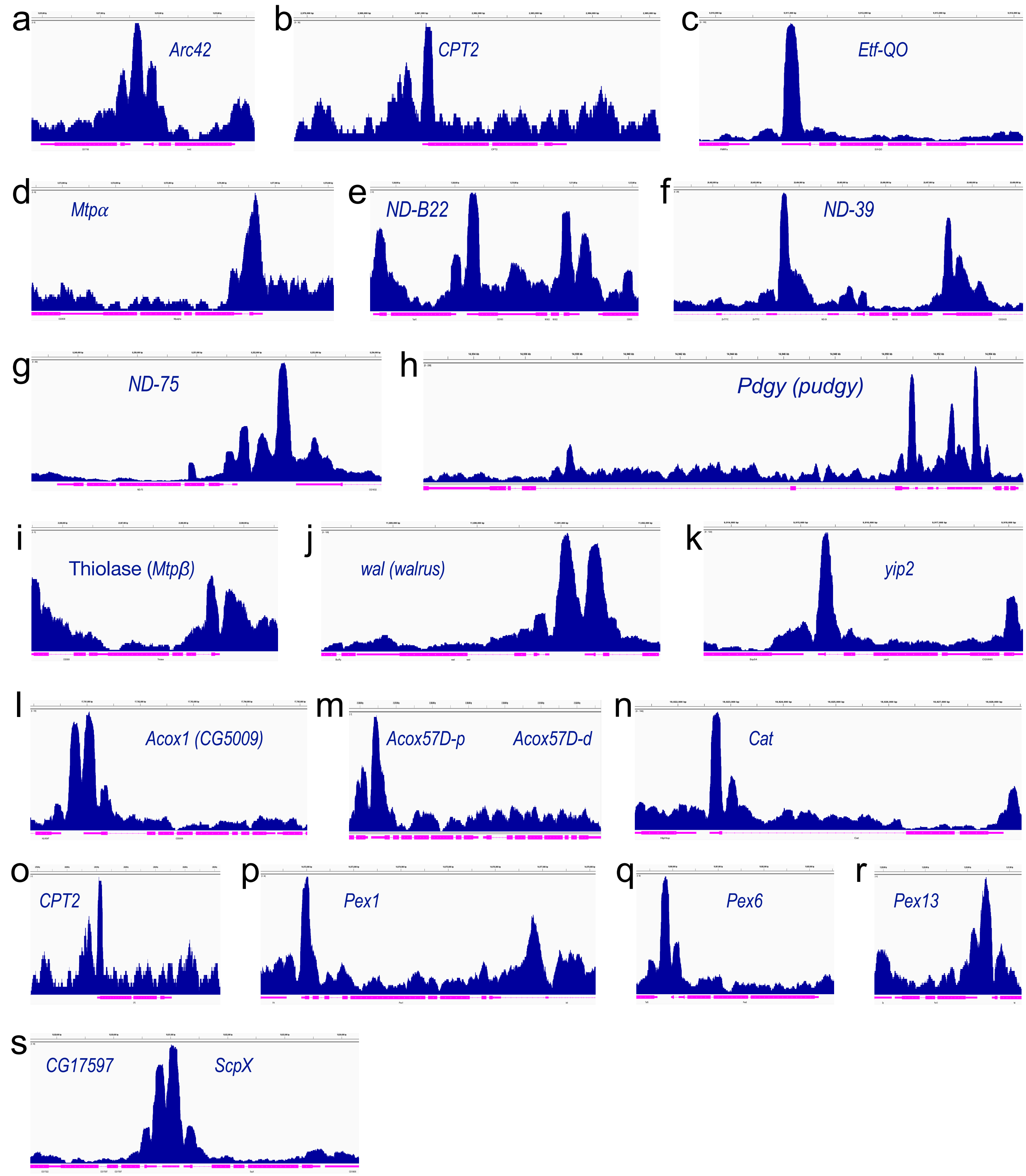

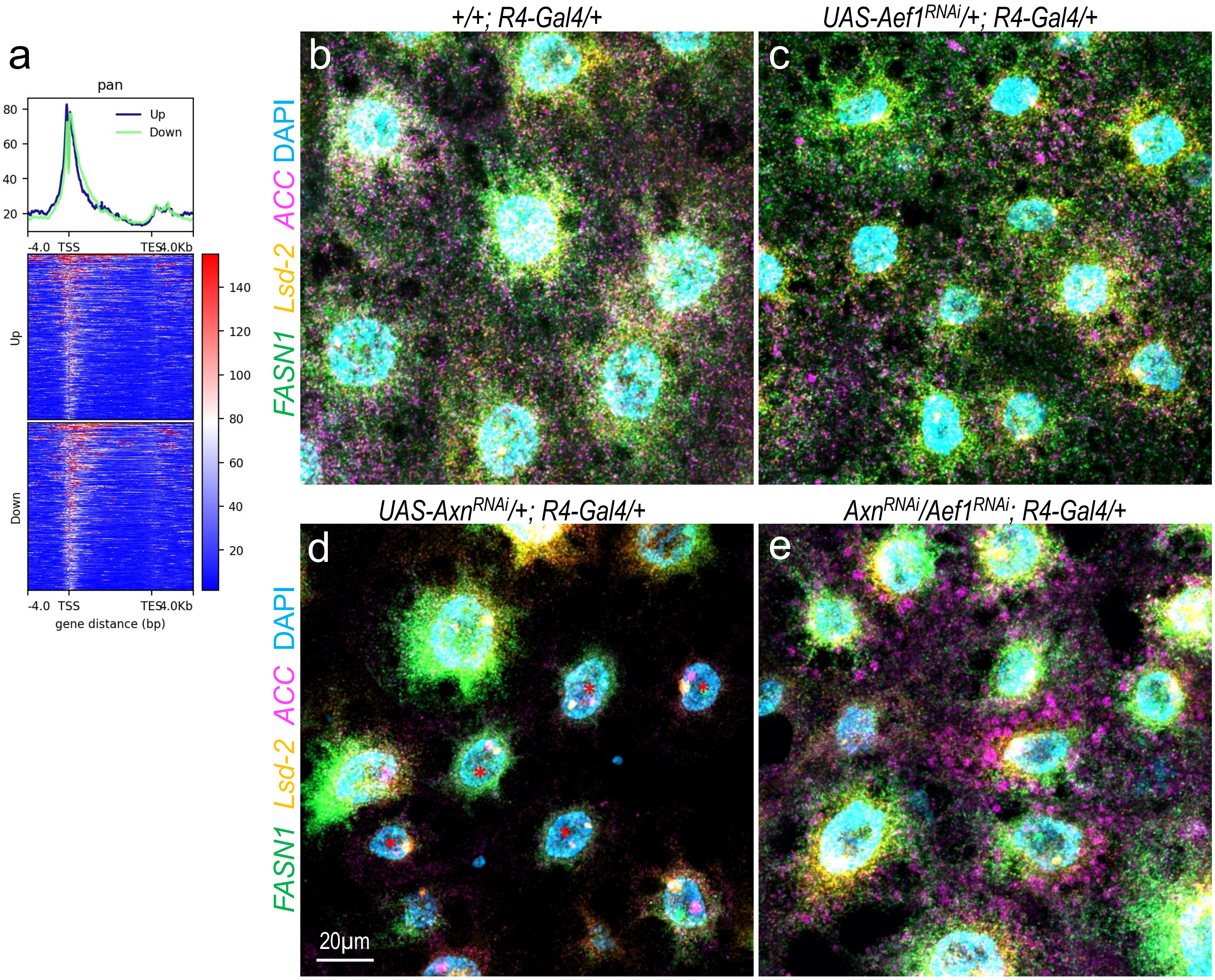


Suppl. Fig. S10
