## Supplementary material for "Wingless signaling promotes lipid mobilization through signal-induced transcriptional repression": Table S1 List of Stocks and Primers

**Table S1 List of the *Drosophila* stocks used in this study**

| Stock # | Genotype | Comments |
|---|---|---|
| 7011 | *w[1118]; P{w[+mC]=Cg-GAL4.A}2* | *dCg-Gal4* |
| 31705 | *y1 v1; P{TRiP.HM04012}attP2* | *UAS-Axn[RNAi]* |
| 33832 | *y[1] w[*]; P{w[+mC]=r4-GAL4}3* | *R4-Gal4 (III)* |
| 33986 | *y1 sc* v1 sev21; P{TRiP.HMS00946}attP2* | *UAS-slmb[RNAi]* |
| 35004 | *y1 sc* v1 sev21; P{TRiP.HMS01414}attP2* | *UAS-arm[RNAi]* |
| 38394 | *y[1] w[*]; wg[Sp-1]/CyO; P{w[+mC]=GAL4-dSREBPg.K}A45/TM6B, Tb[+]* | *SRE-Gal4 (III),* Used with classical genetic recombinations |
| 38395 | *y[1] w[*]; P{w[+mC]=GAL4-dSREBPg.K}A39; SREBP[189]/TM6B, Tb[1]* | *SRE-Gal4 (II),* Used with classical genetic recombinations |
| 40848 | *y1 v1; P{TRiP.HMS02015}attP40/CyO* | *UAS-pan[RNAi]*, Homozygous animals were used |
| 58787 | *w*; P{UAS-α-Cat.T:GFP.sg}3/CyO* | *UAS-α-Cat+* |
| 59759 | *y1 w*; Mi{PT-GFSTF.1}nkdMI00209-GFSTF.1/TM3, Sb1Ser1* | *nkd[EGFP]* |
| 60561 | *y1 w* Mi{PT-GFSTF.1}armMI08675-GFSTF.1* | *arm[EGFP]* |
| 62434 | *y1 v1; P{TRiP.HMJ23888}attP40/CyO* | *UAS-Axn[RNAi]*, Balancer changed to Cyo,Tb |
| 65020 | *y1 sc* v1 sev21; P{TRiP.HMC05894}attP40* | *UAS-Lsd-1[RNAi]* |
| | *Sp/Cyo; fz3-RFP/TM6B* | a kind gift from Dr. Yashi Ahmed |
| | *UAS-Lsd1[EGFP]/TM3,Sb*<br>*UAS-Lsd2[EGFP]/TM3,Sb* | a kind gift from Dr. Mathias Beller |
