## Supplementary material for "Wingless signaling promotes lipid mobilization through signal-induced transcriptional repression": Table S2 Primers used

Table S2 Primers used to generate the *pan*$^{EGFP}$ strain using the CRISPR-Cas9 technique

| Primer name | Primer Sequence | |
|---|---|---|
| 1kb_UPS_Pan 5.1 | ggcggccgcgggaattcgatAAGACGTTTATCAAACATGTTC | |
| 1kb_UPS_Pan 3.1 | aggatctcagATATAAGAGTTACAGAACAGATC | |
| Last_IN_EX_Pan 5.1 | actcttatatCTGAGATCCTTGCTGCATG | |
| Last_IN_EX_Pan 3.1 | tgctcaccatTGAAACGCTAATAACGCC | |
| eGFP_Pan 5.1 | tagcgtttcaATGGTGAGCAAGGGCGAG | *Mutated PAM sites |
| eGFP_Pan 3.1 | tggcgatcagTTACTTGTACAGCTCGTCCATG | |
| 3(pr)_UTR_Pan 5.1 | gtacaagtaaCTGATCGCCATGGATTTG | |
| 3(pr)_UTR_Pan 3.1 | aattaatggaTTTTGGCAAGTTGTGTCTAATTTTAAAATAAAATAC | |
| 1kb_DWNS_Pan 5.1 | cttgccaaaaTCCATTAATTAATGCCTCTCTATCACATATG | |
| 1kb_DWNS_Pan 3.1 | gccgcgaattcactagtgatTACATGGAATAAAGGCTATC | |
| SgRNA1_Pan 5.1 | gtcGATCTGTTCTGTAACTCTTA | |
| SgRNA1_Pan 3.1 | aaacTAAGAGTTACAGAACAGATC | |
| SgRNA2_Pan 5.1 | gtcGTCCATTAATTAATGCCTCTC | |
| SgRNA2_Pan 3.1 | aaacGAGAGGCATTAATTAATGGA | |
| 1kb_UPS_Pan(Seq) 5.1 | AAGACGTTTATCAAACATGTTC | |
| 1kb_UPS_Pan(Seq) 3.1 | ATATAAGAGTTACAGAACAGATC | |
| Last_IN_EX_Pan(Seq) 5.1 | CTGAGATCCTTGCTGCATG | |
| Last_IN_EX_Pan(Seq) 3.1 | TGAAACGCTAATAACGCC | |
| eGFP_Pan(Seq) 5.1 | ATGGTGAGCAAGGGCGAG | |
| eGFP_Pan(Seq) 3.1 | TTACTTGTACAGCTCGTCCATG | |
| 3(pr)_UTR_Pan(Seq) 5.1 | CTGATCGCCATGGATTTG | |
| 3(pr)_UTR_Pan(Seq) 3.1 | TTTTGGCAAGTTGTGTCTAATTTTAAAATAAAATAC | |
| 1kb_DWNS_Pan(Seq) 5.1 | TCCATTAATTAATGCCTCTCTATCACATATG | |
| 1kb_DWNS_Pan(Seq) 3.1 | TACATGGAATAAAGGCTATC | |
| 1kb_UPS(Mid)_Pan(Seq) 5.1 | GTATGTGTCGCGGCAGAGCG | |
| Last_IN_EX(Mid)_Pan(Seq) 5.1 | CTATCATGCTAATCACTCGC | |
